# Single Nuclei-Derived Molecular Subtypes of Gastrointestinal Stromal Tumors Correlate with Clinicopathologic Features and Predict Clinical Outcomes

**DOI:** 10.1101/2025.04.21.647322

**Authors:** Ashwyn K. Sharma, Alexander T. Wenzel, John Jun, Diwakar Guragain, Chih-Min Tang, Mark Antkowiak, Shruti Bhargava, Adam M. Burgoyne, Paul Fanta, Sadie E. Munter, Katherine Y. Chen, Terence M. Doherty, Manikandan Murugesan, Adrian Sanchez-Fdez, Shashwath Malli, Prabhjot S. Mundi, Sara Rothschild, Denisse Evans, William Kim, Gary K. Schwartz, Andrea Califano, Pablo Tamayo, Jill P. Mesirov, Jason K. Sicklick

**Affiliations:** Department of Surgery, Division of Surgical Oncology, University of California San Diego, San Diego, CA, USA; Moores Cancer Center, University of California San Diego, La Jolla, CA, USA; Department of Medicine, Division of Genomics and Precision Medicine, University of California San Diego, San Diego, CA, USA; Department of Pharmacology, University of California San Diego, San Diego, CA, USA; Department of Medicine, Vagelos College of Physicians and Surgeons, Columbia University, New York, NY, USA; Herbert Irving Comprehensive Cancer Center, Columbia University, New York, NY, USA; The Life Raft Group, Wayne, NJ 07470; Case Comprehensive Cancer Center, Cleveland, OH 44106, USA; University Hospitals Cleveland Medical Center, Cleveland, Ohio; Department of Systems Biology, Columbia University, New York, NY, USA; Department of Biochemistry and Molecular Biophysics, Vagelos College of Physicians and Surgeons, Columbia University, New York, NY, USA; Department of Biomedical Informatics, Vagelos College of Physicians and Surgeons, Columbia University, New York, NY, USA; Chan Zuckerberg Biohub New York, New York, NY, USA

## Abstract

Historically, gastrointestinal stromal tumor (GIST) has been subtyped by oncogenic driver mutations. However, tumors with the same mutational profile can have variable biology. To further explore the impact of molecular diversity on GIST biology, we performed single nucleus RNA sequencing on 16 primary GIST and utilized an integrated single cell atlas of the normal GI tract composed from multiple publicly available datasets to identify six distinct GIST *cell states*. We then statistically estimated the relative abundances of these profiles in bulk transcriptomic data. These were used to define six common GIST *molecular subtypes* based upon one or two predominant tumor *cell states*. We found that these *molecular subtypes* correlate with tumor locations, mutational profiles, and patient outcomes, and validated these subtypes in an independent international cohort. These *molecular subtypes* have the potential to be used for clinical prognostication for patients with GIST, identifying new therapeutic targets, and studying the cell of transformation of GIST.

## INTRODUCTION

Gastrointestinal Stromal Tumor (GIST) constitutes a genomically diverse mesenchymal cancer that arises throughout the gut. Approximately 60–70% of GISTs are driven by gain-of-function mutations in *KIT*. Another 20–25% are caused by oncogenic mutations in *PDGFRA* and *BRAF,* or *ETV6-NTRK3* and *FGFR1* fusions^1–9^. Additional subtypes are caused by either germline or somatic mutations in the neurofibromatosis type 1 (*NF1*) gene, as well as germline mutations in succinate dehydrogenase (*SDH*) subunit genes ^10,11^.

Interestingly, the mutational landscape of GIST also correlates with tumor locations ^12–14^. In fact, *PDGFRA* and *SDH* mutant GIST solely arise in the stomach, whereas *BRAF^V600E^* mutant and somatic *NF1* mutant GIST commonly arise at the duodenal-jejunal flexure^15^. Furthermore, even within the stomach, there are spatial predilections^16^. Recent evidence from our group has demonstrated that proximal gastric GIST are 96% *KIT* mutant, while distal gastric tumors are 74.4% non-*KIT* mutant^16^. These findings raise biological questions regarding the tumoral heterogeneity of GIST.

To date, there is limited direct evidence that oncogenic mutations in a specific cell type can cause GIST tumorigenesis. Conventionally, GIST has been believed to arise from ANO1 (DOG-1) and CD117 (KIT) expressing interstitial cells of Cajal (ICCs), which function as myo-fibroblastic pacemaker cells of the GI tract. It was reported that myenteric and intra-muscular ICCs may be transformed by a *KIT* mutation in a genetically engineered mouse model (GEMM)^17^. However, other cell types now have been implicated in GIST tumorigenesis. Recently, telocytes, fibroblast-like cells expressing PDGFRA and CD34, were postulated as the ICC equivalent in one patient with germline *PDGFRA* mutant GIST ^18^. Moreover, smooth muscle cells of the small intestine have been shown to give rise to *BRAF*^V600E^ mutant GIST in a different GEMM ^19^.

Given the genomic-spatial correlations in GIST and the different postulated benign precursors of GIST described above, we hypothesized that: 1) GIST may exhibit heterogeneous transcriptional profiles; 2) that these profiles may vary by tumor location and/or driver mutation; and 3) that these profiles may be prognostic of patient outcomes. Herein, we compose a single cell atlas of the human mesenchymal GI tract by performing a pooled analysis of publicly available datasets and used this to investigate the inter- and intra-tumoral heterogeneity of GIST at a single cell resolution.

## RESULTS

### Single cell reference atlas of normal mesenchymal and enteric neural cells

We utilized a multi-step workflow for our analysis, in which we first sought to examine the normal GI tract at a single cell resolution (**Figure 1**). To do this, we utilized 21 publicly available single cell RNA sequencing (scRNAseq) and single nuclei RNA sequencing (snRNAseq) datasets of the human gastrointestinal tract that had been previously annotated via marker genes and literature review^20–39^. We then integrated cells of mesenchymal origin and the enteric nervous system (ENS) while excluding immune, epithelial, and other cell populations to create a reference atlas of the normal mesenchymal and enteric neural cells of the GI tract (see **Methods**). Overall, the combined reference dataset encompassed more than 300,000 cells or nuclei from more than 700 samples from unique patients, as well as spanned all portions of the GI tract (i.e., esophagus to rectum) across the lifespan (i.e., fetal development to adulthood).

**FIGURE 1.**
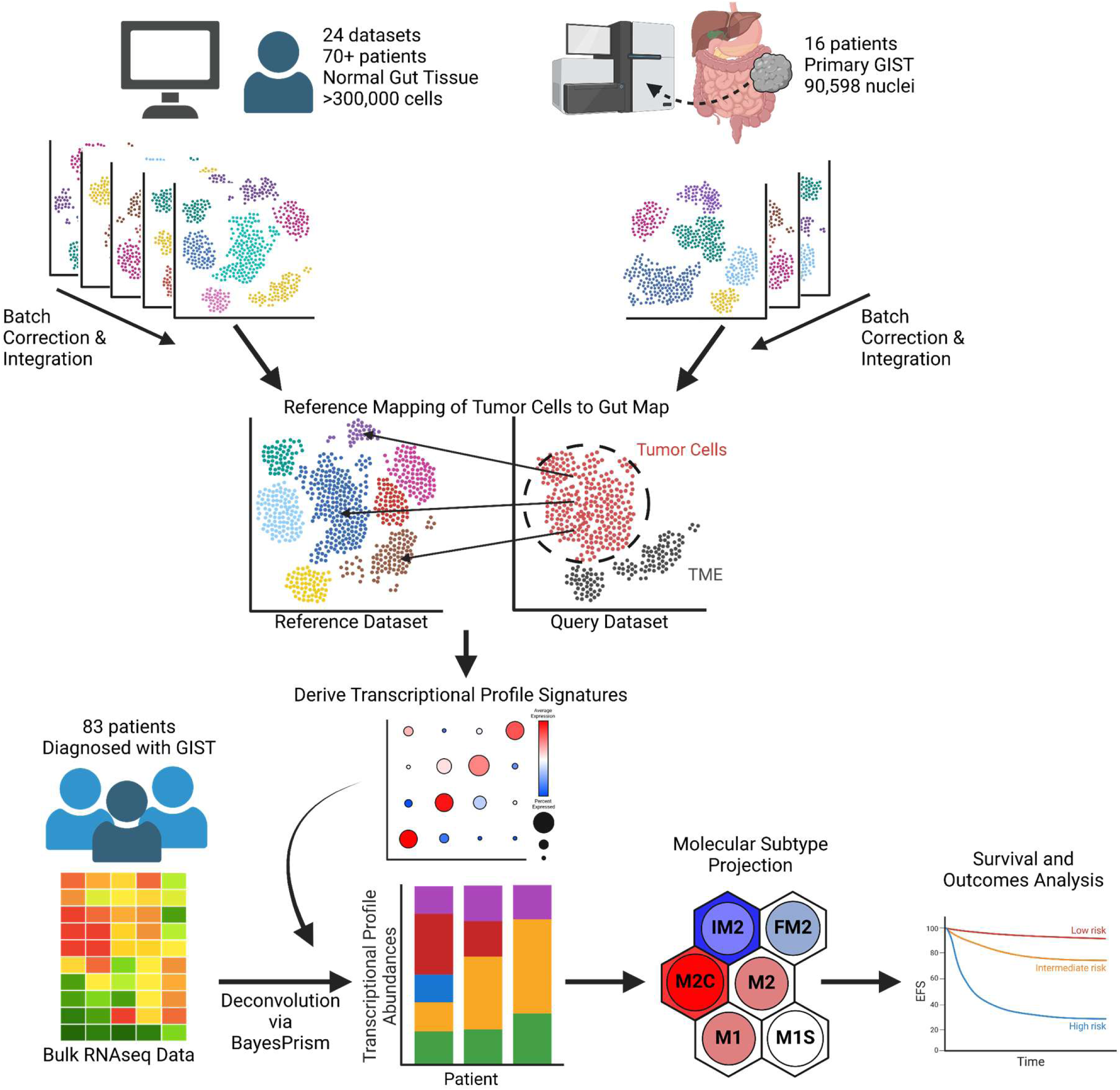
Schematic Workflow demonstrating Bioinformatic Analysis. First, publicly available datasets were batch corrected and integrated to create a reference dataset of the mesenchymal and enteric neural human GI tract. In addition, snRNA sequencing was concurrently performed, batch corrected, and integrated to identify all GIST populations. Using referencing mapping, tumor cells were projected onto our reference map to identify *cell states* and their respective transcriptional profiles. Using these profiles, we performed deconvolution of bulk RNAseq data from 83 patients to derive *cell state* abundances for each sample. We then used a Self-Organizing Map clustering algorithm, using the *cell state* abundances as input, to derive *molecular subtypes* for these samples. *Molecular subtypes* correlated with clinical and survival outcomes.

Given the significant diversity and granularity of cell types in this integrated normal atlas, we attempted to broadly categorize these cells. Cell clustering by a shared nearest neighbor (SNN) modularity optimization-based clustering algorithm ^40,41^ demonstrated the presence of multiple large groups of cells (**Figure 2A**). These were then annotated based upon a combination of labels assigned in the original study from which a cell originated, as well as calculated marker genes **(Supplementary Figure 1A**). Fibroblasts and stromal cells clustered together, as did smooth muscle cells. Neuronal cells and enteric glia clustered separately from these populations, and consisted of cells such as motor neurons, neuroblasts, and glia. Two fetal mesodermal populations, Mesoderm_1, characterized by *HAND1*, and Mesoderm_2, characterized by *ZEB2, SHISHA3 and TCF21* expression, were also identified. Notably, expression of *HAND1* has been associated with small bowel GIST location and more aggressive tumor biology^42^. Meanwhile, the Mesoderm 2 population, characterized by *ZEB2* expression, has been recently demonstrated to promote the development of neural lineages in murine models^43^. We also noted a population of cycling stromal cells characterized by cell cycle genes such as *MIK67*, *TOP2A*, and *CDK1*. Finally, ICCs were identified by their expected markers, *KIT*, *ANO1*, and *ETV1*.

**FIGURE 2.**
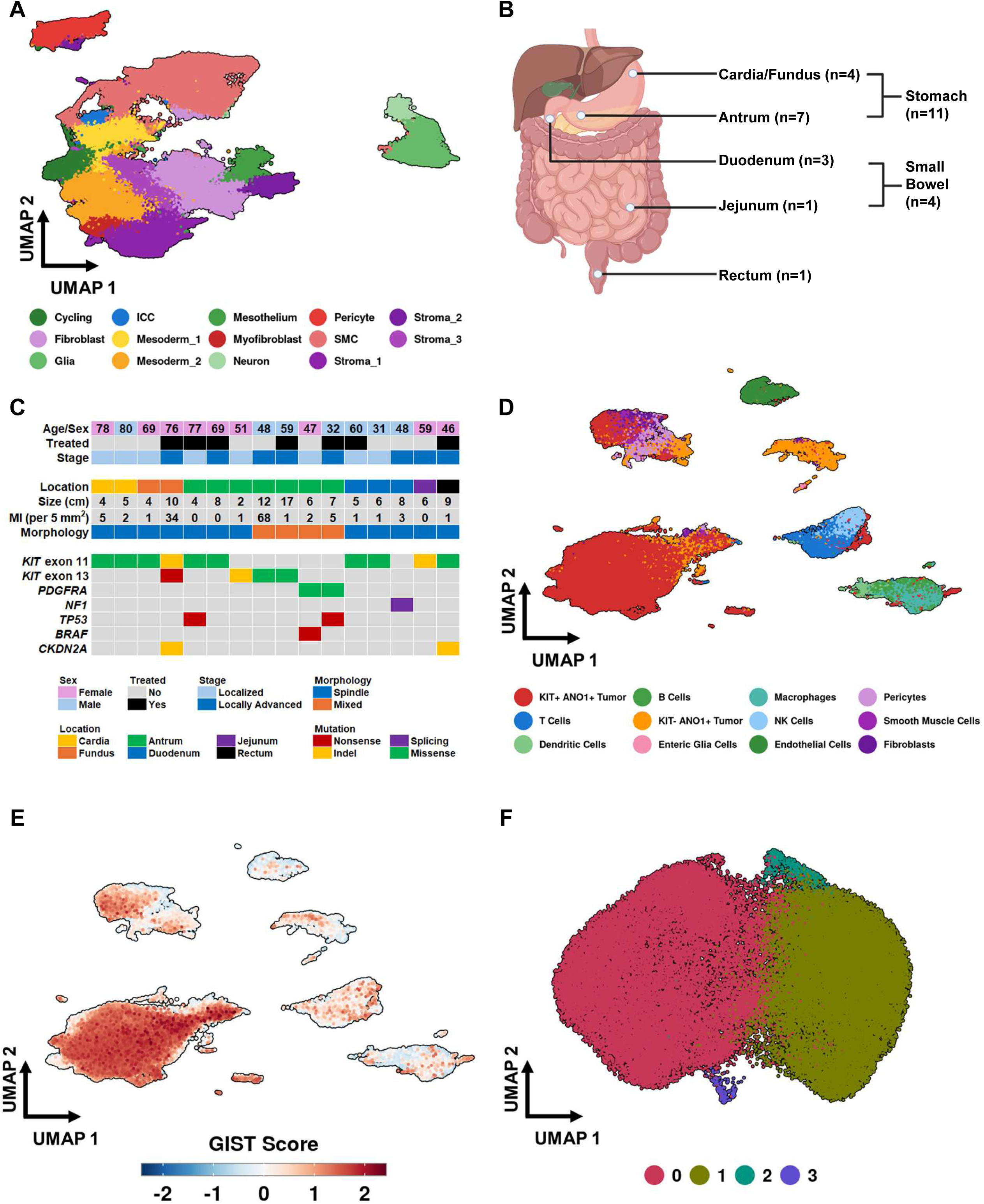
Human Gut Reference Map and the snRNAseq Atlas of Human Primary GIST. **A.** Uniform Manifold Approximation and Projection (UMAP) of normal GI tissue procured from publicly available datasets. Cells were categorized into major groups based on marker genes and broad characterization. **B.** Sample distribution of 16 tumors by GI location. **C.** Clinical and demographic information for the 16 samples chosen to undergo snRNA sequencing. Patients were selected based on differing mutation status, location, mitotic rate, gender, and age. **D.** UMAP of 16 human GISTs and the tumor microenvironment, with multiple tumor cell populations labeled. **E.** Feature Plot of GIST scores, a composite expression measure of the following genes: *KIT, ANO1, ETV1, PRKCQ, GPR20, PDE1A, PROM1,* and *FOXF1,* for each human GIST tumor cell. Red indicates relative high scores of these markers. **F**. UMAP of all *KIT*^+^ and *KIT^-^* tumor cells.

### Single nuclei RNA sequencing of human primary GIST illustrates tumor heterogeneity

Next, we performed snRNAseq of primary human GIST from 16 different patients to characterize inter- and intra-tumoral heterogeneity. These samples encompassed 3 general locations (stomach, n=11; small bowel, n=4; rectum, n=1) (**Figure 2B**) and multiple driver mutations (*KIT*, n=10; *PDGFRA*, n=3; *SDH*, n=2; *NF1*, n=1). We included patients across the age spectrum (range: 31-80 years old), as well as both untreated and treated tumor samples (**Figure 2C**). After quality filtering, 90,598 nuclei from the 16 samples were included for further analysis.

Using markers such as *ANO1*, *KIT* and *ETV1*, we identified cell clusters that were likely tumor cells. Consistent with previous studies, we identified two major tumor populations: a *KIT*^+^ *ETV1*^+^ *ANO1^+^* tumor population, and a *KIT^-^ ETV1^-^ ANO1^+^* tumor population^44^ (**Figure 2D**). Given the non-specificity of *ANO1* alone as a marker for GIST, we then compiled a composite GIST expression score (see **Methods**) for each individual cell based on expression of other known GIST markers, including *PRKCQ*, *PDE1A*, *PROM1* (CD133), and *FOXF1* (**Figure 2E**). The *KIT*^+^ *ETV1*^+^ *ANO1^+^* and *KIT^-^ ETV1^-^ ANO1^+^*tumor populations both scored high for GIST markers in comparison to other cell populations, indicating that these populations likely reflect true tumor cell populations. Within our samples, other identified cell types included immune cells (B, T, NK, and dendritic cells), epithelial cells, stromal cells (including fibroblasts), smooth muscle cells, pericytes, endothelial cells, and enteric glial cells.

Sub-clustering of all tumor cells (i.e., pooled *KIT*^+^ and *KIT*^-^ tumor cells) demonstrated 4 distinct clusters (**Figure 2F, Supplementary Figure 1B**). Recognizing the importance of treatment status, we also explored the differences in gene expression between treated and untreated tumor cells. Treated tumor cells had higher expression in genes such as *DGKH* and *HMCN1* **(Supplementary Figure 1C**). But expression of GIST markers such as *KIT*, *ANO1*, and *ETV1* were similar between treated and untreated tumor cells **(Supplementary Figure 1D**).

### Human GIST demonstrates transcriptional similarity to multiple mesenchymal cell types other than ICCs

Given the genomic and biological diversity of GIST, as well as the presence of multiple tumor sub-clusters in the aforementioned analysis, we next hypothesized that GIST may have different *cell states* which are characterized by specific transcriptional profiles. To further investigate this, we performed reference mapping of tumor cells onto the integrated human gut single cell reference map described above. First, we performed a validation of our methodology of combining snRNAseq and scRNAseq data by mapping snRNAseq tumor cells onto a previously published scRNAseq dataset consisting of two GISTs^45^ **(Supplemental Figure 2A-B**). We then projected tumor cells onto the reference map using the Symphony package^46^ and assigned a label based on transcriptional similarity (i.e., a tumor cell that projects to ICCs is labeled as an ICC). This validation demonstrated appropriate labeling and high concordance of matching cell types (reference matching similarity scores were between 0.8 and 1.0, data not shown).

Consistent with the hypothesized benign precursor of GIST, 62.3% (n=38,008) of tumor cells projected onto ICCs (**Figure 3A-B**) in the normal gut reference map (in grey). However, 37.7% of tumor cells projected to other cell types, including Smooth Muscle Cells (SMC; n=10,706; 17.6%), Fibroblasts (n=5,889; 9.8%) Mesoderm_1 (n=5,831, 9.6%), Mesoderm_2 (n=367, 0.6%), and Cycling cells (n=73, 0.1%). Since these single nuclei represent tumor cells, we elected to refer to these differentially mapped subpopulations as distinct GIST tumor *cell states*. Cell state distributions for each sample from our projection can be found in **Supplemental Figure 2C**.

**FIGURE 3.**
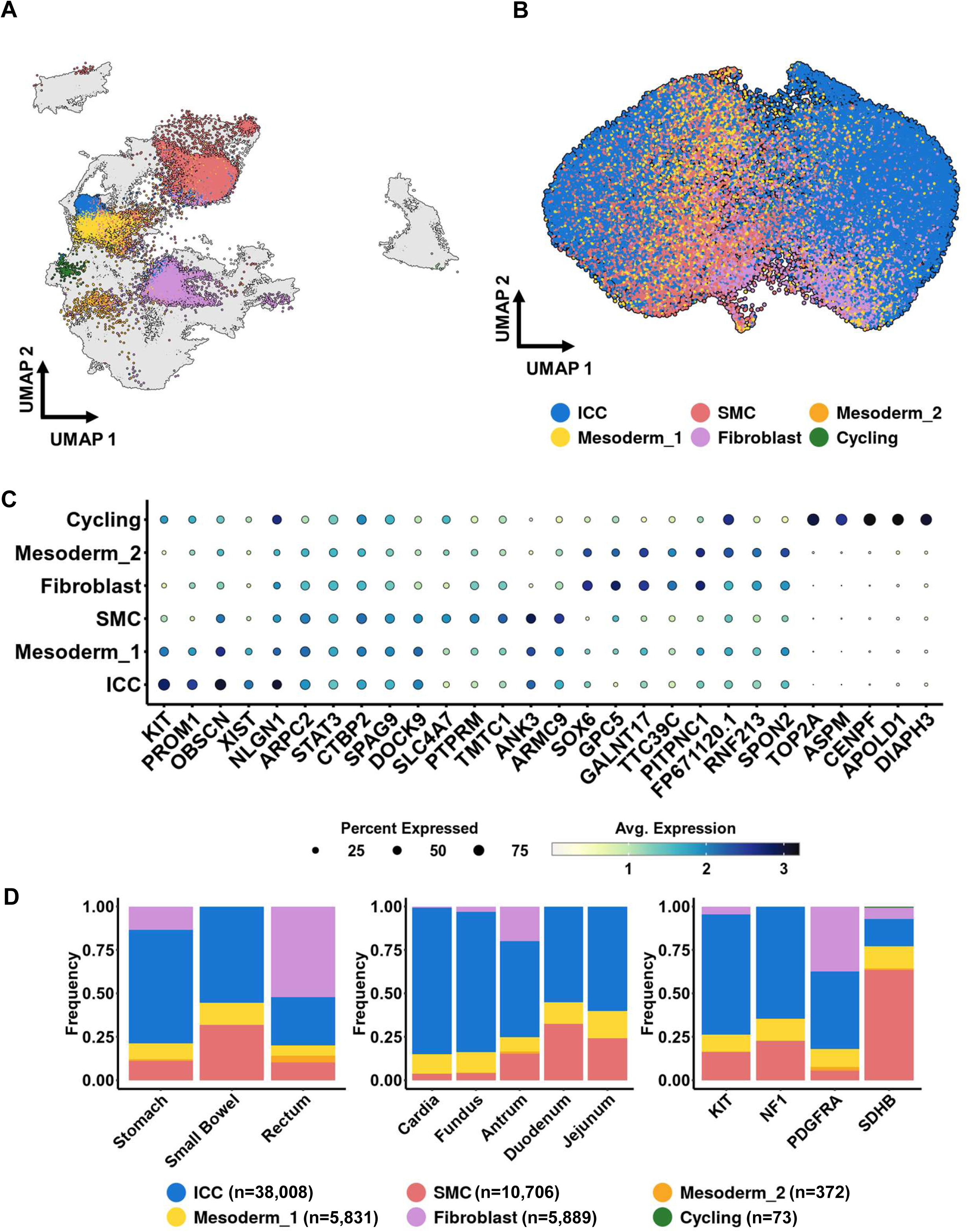
Identification of ICC-like and non-ICC like Tumor Cell States by Reference Mapping. **A.** Tumor Cells (colored) are projected onto Gut Reference Map (grey scale). Projection results demonstrate that nearly two-thirds of tumor cells map to ICCs while the remaining cells map to other *cell states*. Blue = ICC, Light Yellow = Mesoderm_1, Yellow-Orange: Mesoderm_2, Purple: Fibroblast, Red: SMC, Green: Cycling. **B.** UMAP of *KIT^+^*and *KIT^-^* tumor cells colored by assigned *cell state*. Blue = ICC, Light Yellow = Mesoderm_1, Yellow-Orange: Mesoderm_2, Purple: Fibroblast, Red: SMC. **C.** Marker Genes of assigned *cell states*. Marker genes were calculated by differential expression using a Wilcoxon rank sum test, with the top 5 genes per cluster shown for brevity. Dot size indicates percentage of cells in each group expressing the gene of interest, and the color indicates average expression level. **D.** Left: Mapped tumor cell types by general location (stomach, small bowel, rectum). Middle: Mapped tumor cell types by specific location, including localized areas of the stomach (cardia, fundus, antrum) and small bowel (duodenum, jejunum). Right: Mapped tumor cell types by driver mutation. Each bar graph represents what proportion of cells in each location or mutation sub-group were assigned a particular *cell state*.

The top marker genes were calculated for each tumor *cell state* based on differential gene expression (**Figure 3C**). The ICC *cell state* was characterized by *KIT* expression (average log_2_FC: 2.04, p < 0.001). The Mesoderm_1 tumor *cell sta*te had higher expression of *SPAG9* (average log_2_FC: 0.441, p < 0.001), while the Fibroblast tumor *cell states* had unique expression of *GPC5* and *PITPNC6*. Cycling cells were characterized by genes involved in DNA repair and assembly such as *TOP2A*, *APOLD1*, and *CENPF*.

We then examined the association between tumor *cell states* and clinical metadata such as primary tumor location and canonical mutation status (**Figure 3D**). Foregut GIST within the proximal stomach (i.e., cardia and fundus) had predominantly ICC-like tumor cells, whereas tumors in the distal stomach (i.e., antrum) were more likely to have non-ICC states (fundus, 19.1% non-ICC; cardia, 15.8% non-ICC vs. antrum, 45.7% non-ICC; p < 0.001), including Fibroblast-like and SMC-like states, paralleling our earlier finding that tumor mutation status correlates with primary tumor location^16^ (**Figure 3D, middle panel)**. In general, more distal GISTs of the embryologic foregut and midgut tended to have lower proportions of ICC-like tumor cells (linear model β = −0.1, p = 0.0146). We next assessed the tumor *cell states* relative to tumor mutation status (**Figure 3D, right panel)**. Tumors with non-*KIT* mutations were found to have more tumor cells possessing non-ICC-like *cell states* when compared to those with *KIT* mutations. For example, *SDHB* mutant tumors were higher in SMC-like states (*SDH*, 63.4% vs *KIT*, 16.2%; p < 0.001), whereas *PDGFRA* mutant tumors were higher in fibroblast-like states (*PDGFRA,* 37.3% vs *KIT,* 4.6%; p < 0.001). *NF1* mutant tumor cells appeared most like *KIT* mutant tumors, which were predominantly comprised of the ICC-like *cell state*. Finally, we examined the association between treatment status and tumor *cell states*. Cycling cells were more likely to be treated cells (p < 0.001, data not shown), although it remains to be determined if this is an effect of treatment and/or indicative of more aggressive tumors that were treated. In contrast, there were no significant differences in distributions of other tumor *cell states* in treated versus untreated cells (**Supplementary Figure 2D**). This suggests that treatment alone does not explain the tumor cell heterogeneity.

### GIST Evolution in response to treatment

Given that our cohort included both untreated and treated tumor cells, we sought to better understand how treatment may transiently influence cellular states using *in vitro* models. To do this, we performed single cell RNAseq (scRNAseq) on *KIT* exon 11 mutant (Δ560-578) GIST-T1 cell lines that were either treatment naïve (T1), treated with imatinib (T1-IM), or resistant to imatinib (T1-IM^R^) (**Figure 4A).** As described above, we once again projected our dataset onto our previously defined reference map (**Figure 4B**). Most cells displayed an ICC cell state (n=27,540, 42.6%) or Mesoderm_1 cell state (n=25,356, 39.2%) (**Figures 4C-D**).

**FIGURE 4.**
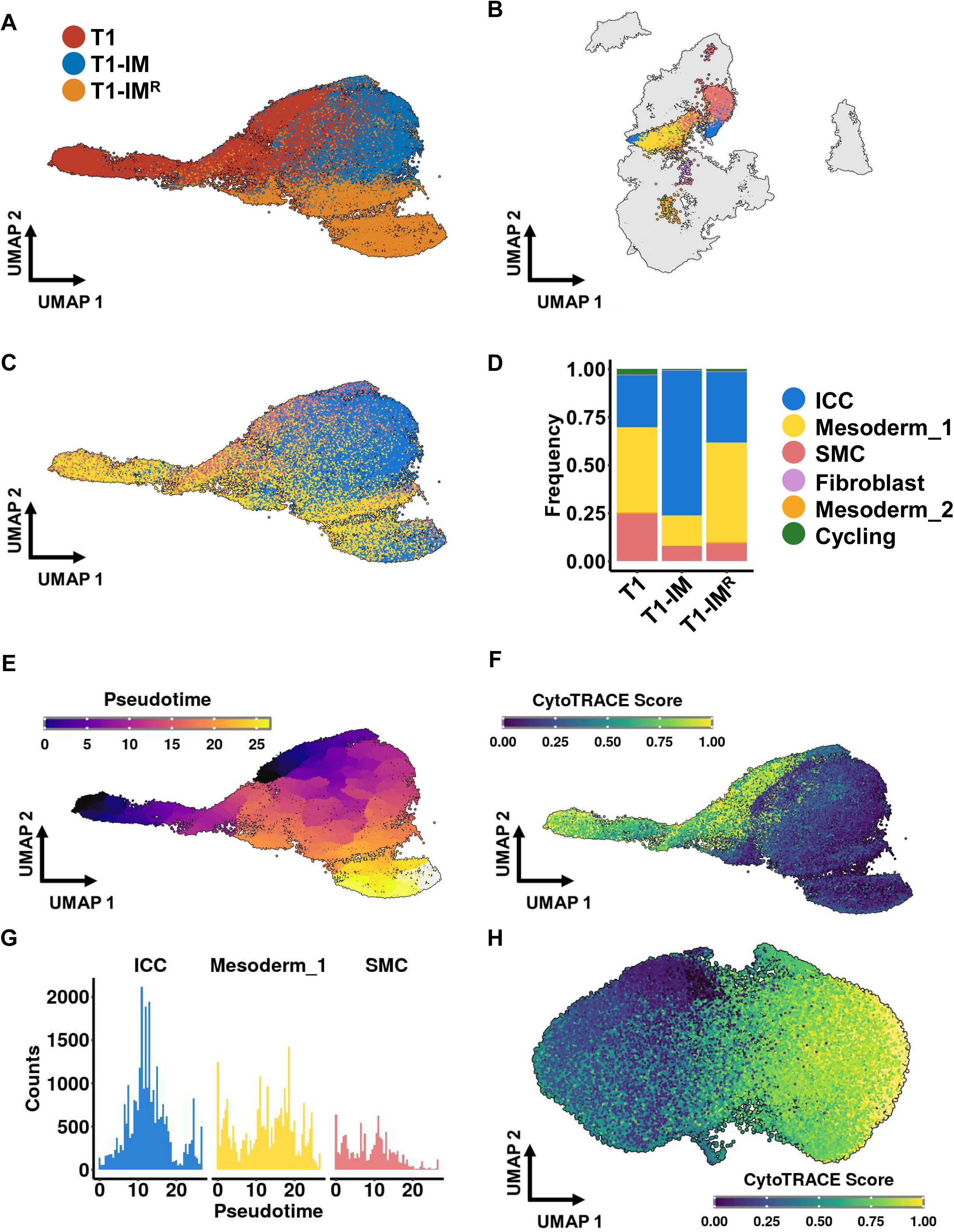
Evolution in Tumor Cell States in Pseudotime. **A.** UMAP of single cell RNA sequencing results of three GIST cell lines, T1 untreated cell lines (red), T1 cells treated with imatinib (blue, T1-IM), and T1 cells resistant to imatinib (orange, T1-IM^R^). **B.** Projection results of three GIST cell lines onto previous reference map. Blue = ICC, Light Yellow = Mesoderm_1, Yellow-Orange: Mesoderm_2, Purple: Fibroblast, Red: SMC. **C.** UMAP of three GIST cell lines based on assigned cell state. Blue = ICC, Light Yellow = Mesoderm_1, Yellow-Orange: Mesoderm_2, Purple: Fibroblast, Red: SMC. **D.** Proportions (range 0.00 – 1.00) of cell states within each different cell line. **E.** UMAP of 3 GIST cell lines based on pseudotime assigned by Monocle. T1 untreated cells were selected as the initialization point for pseudotime analysis. **F.** UMAP of 3 GIST cell lines based on pseudotime assigned by CytoTRACE (CT). A CT score of 0.00 is considered later in pseudotime/more differentiated, while a CT score of 1.00 is considered earlier in pseudotime and more “undifferentiated”. **G.** Histogram demonstrating changes in the number of cells in a particular cell state over pseudotime. The three most common cell states (ICC, Mesoderm_1, and SMC) are displayed, other cell states were omitted due to low number of cells assigned to these states. **H.** UMAP of all human *KIT*^+^ and *KIT^-^*tumor cells with assigned CT scores. A CT score of 0.00 is considered later in pseudotime/more differentiated, while a CT score of 1.00 is considered earlier in pseudotime and more “undifferentiated”.

To quantify how tumor cells change in response to treatment, we performed pseudotime analysis using the Monocle package, which strives to order cells based on gene expression into cell trajectories^47–49^. Using untreated cells as an initialization point (pseudotime = 0), tumor cells are ordered over pseudotime. As expected, over pseudotime, T1 cells in response to therapy follow a progression towards T1-IM cells and then proceed towards T1-IM^R^ cells (**Figure 4E**). However, because Monocle requires prior knowledge of the initial starting point to perform pseudotime analysis, we confirmed our findings using a second pseudotime analysis package,

CytoTRACE (CT), which orders cells in pseudotime using gene counts and expression without the need for setting an initialization point, with “de-differentiated” or cells ordered earlier in pseudotime having higher CT scores^23^. This yielded T1 cells with higher CT scores, whereas T1-IM and T1-IM^R^ populations had lower CT scores (**Figure 4F**). When examining changes in cellular state over pseudotime, we noted that treatment is associated with a transient change in cell state, with an increase in the ICC cell state (**Figure 4G, Supplemental Figure 3)**. Over pseudotime, we also note an enrichment in the Mesoderm_1 cell state by the time T1 cells become resistant to imatinib. Taken together, our snRNAseq analyses of human tumors and scRNAseq analyses of the GIST-T1 cell line demonstrate that GIST have inter-tumoral and intra-tumor heterogeneity that evolves with treatment. Finally, using CytoTRACE (CT) on human snRNAseq data demonstrates a progression of cells in pseudotime (**Figure 4H**). While snRNAseq and scRNAseq can be utilized for research questions, their potential application for clinical application is limited at present. Therefore, with our understanding of tumor cell states in GIST, we sought to investigate whether this heterogeneity could be appreciated using bulk RNAseq data from human tumors.

### Molecular Subtypes of Clinical GIST correlate with survival outcomes

We collected bulk RNAseq data from 83 patients who had been diagnosed with GIST and had undergone Tempus Next Generation Sequencing (NGS) as part of routine clinical care. Using our single nuclei subtypes as a reference, we used the BayesPrism package^50^, which employs Bayesian models, to deconvolve bulk RNAseq data into *cell state abundances* using each *cell state’s* representative expression profile (**Figure 5A**). Prior to performing bulk RNAseq deconvolution, we validated this method on randomly generated pseudomixtures derived from our snRNAseq data. We noted high concordance (median correlation: 0.87) between the predicted fractions and the actual, generated pseudomixture compositions **(Supplemental Figure 4)**.

**FIGURE 5.**
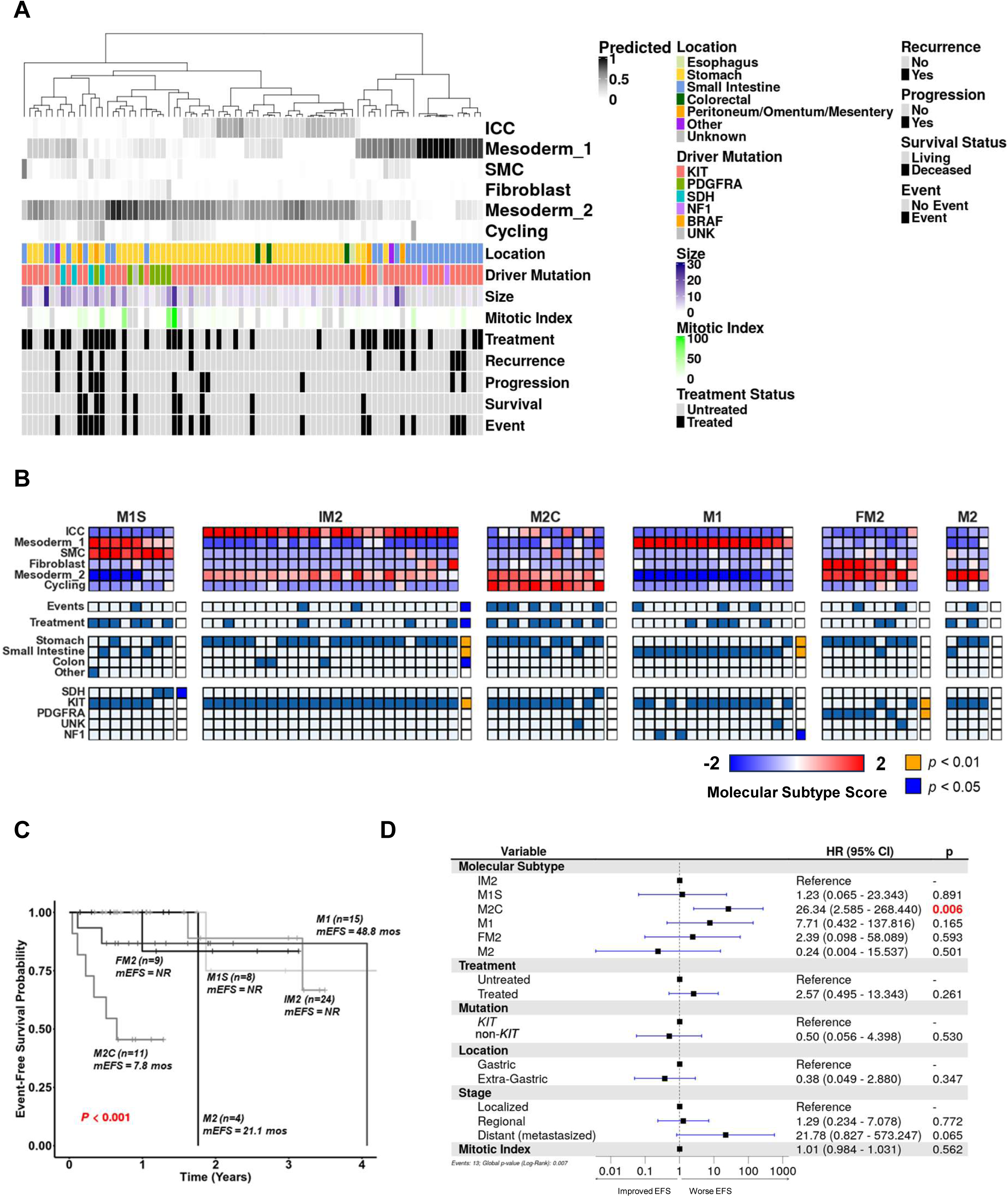
From single cell to bulk RNA sequencing: Deconvolution and Molecular Subtype Projection. **A.** Deconvolution of n=83 bulk RNAseq profiles from patients who underwent tumor biopsy or resection at UC San Diego Health. Proportions of each projected cell-type are determined for each sample. Meta-data is displayed for each sample. **B.** *Molecular subtype* projection of the deconvolution data gives rise to 6 major *molecular subtypes* for tumors. *M1S* = Mesoderm_1 > SMC; 2) *IM2* = ICC > Mesoderm_2; 3) *M2C* (Mesoderm_2 > Cycling); 4) *M1* (Mesoderm_1); 5) *FM2* (Fibroblast > Mesoderm_2); and 6) *M2* (Mesoderm_2). Certain *molecular subtypes* are associated with particular clinical metadata (p < 0.05, blue box, p<0.01, gold box). **C.** Univariate Kaplan-Meier Curve of patient event-free survival based upon *molecular subtype*. **D.** Forest plot of Multivariate Cox Proportional Hazard model for event-free survival (EFS) based on commonly accepted prognostic factors as well as *molecular subtype* projections. *Molecular subtypes* were significant predictors of EFS even when controlled for clinical factors.

Most deconvolved bulk RNAseq tumor samples consisted of at least one or two predominant *cell states*. Moreover, these *cell state* combinations were recurring (i.e., non-random). Therefore, we used a Self-Organizing Map (SOM, see **Methods**) clustering algorithm to group our deconvolved samples into defined patterns, which we called *Molecular Subtypes*. These represent the most salient and common combinations of *cell states* (**Figure 5B**). In total, 71 of the 83 samples (86%) closely matched to one of the *molecular subtypes* according to the Information Coefficient (IC), a mutual information measure of correlation similar to the Pearson correlation coefficient that can detect non-linear associations^51–53^. The remaining 12 samples did not match *molecular subtype profiles* closely (i.e., did not meet an IC cutoff of 0.6). Overall, the mean IC for matched samples to their closest *molecular subtype* profile was 0.782 (min: 0.705, max: 0.840) The six *molecular subtypes* were named based on their predominant *cell state* abundances as follows: 1) *M1S* (Mesoderm_1 and SMC); 2) *IM2* (ICC and Mesoderm_2); 3) *M2C* (Mesoderm_2 and Cycling); 4) *M1* (Mesoderm_1); 5) *FM2* (Fibroblast and Mesoderm_2); and 6) *M2* (Mesoderm_2). It is noteworthy that all 6 *molecular subtypes* included fetal mesodermal *cell state* signatures (i.e., Mesoderm_1 in 33% and Mesoderm 2 in 67%).

Previously, expression of the *HAND1* transcription factor (included within the Mesoderm_1 signature) has been shown to correlate with aggressive biology in small bowel GISTs, while expression of the *BARX1* transcription factor was associated with indolent biology in gastric GIST^42^. To investigate this finding further, we sought to determine if our newly identified *molecular subtypes* correlated with driver mutation, location, and outcomes (i.e., recurrence, progression or death) (**Figure 5B**). In fact, the M2 molecular subtypes were associated with gastric GIST. The *IM2 molecular subtype* consisted of primarily *KIT* mutant gastric GIST (p < 0.01) while *PDGFRA* mutant tumors were more likely to be the *FM2 molecular subtype* (p < 0.01) **(Supplemental Figure 5A-B**). Consistent with Hemming *et al.*, small bowel tumors generally tended to consist of an *M1* subtype (p < 0.01) ^42^. In fact, *NF1* mutant GIST, which generally only occur in the small bowel, were exclusively categorized as the *M1* subtype (p < 0.01). We further examined exon data and found that the *IM2 molecular subtype* was most associated with *KIT* exon 11 mutations (p < 0.01), while the *M1 molecular subtype* was most associated with *KIT* exon 9 mutations (p < 0.01) (**Supplemental Figure 6A)**. However, in contrast to the results of Hemming *et al.*, the *M1S molecular subtype* occurred in both the stomach and small bowel. Moreover, *SDHB* mutant tumors of the stomach were more likely to be of the *M1S molecular subtype* (p < 0.05). These findings both confirm and challenge the earlier report that HAND1 expression (i.e., M1) only marks GIST in the small bowel.^42^ We also investigated secondary *KIT* mutations (exons 13, 14, 17, or 18) that drive tyrosine kinase inhibitor (TKI) resistance. They were present in the *M2* (p < 0.05), *M1S* (p > 0.05, not significant), and *M2C* subtypes (p > 0.05, not significant).

Importantly, these *molecular subtypes* correlated with patient outcomes. We tabulated clinicopathologic data including tumor location and mutation status, as well as assessed outcomes using event-free survival (EFS), a composite outcome defined as one or more of the following: 1) recurrence of disease, 2) progression of disease, or 3) death from any cause. Kaplan-Meier analysis demonstrated that the *M2C molecular subtype* was associated with the shortest EFS (median EFS: 7.8 months, p < 0.001, log-rank test) (**Figure 5C**), while the *IM2 molecular subtype* was associated with the longest EFS (median EFS: not reached). Recognizing that secondary *KIT* mutations are an important predictor of drug resistance and poor outcomes, we noted that the *M2C* subtype was independent of the presence of secondary *KIT* mutations (Fischer’s exact test, p = 0.231). We then performed a multivariate Cox Proportional Hazard Ratio model (**Figure 5D**) after generating univariate Kaplan-Meier survival curves based on *molecular subtypes.* We included *molecular subtype* membership, in addition to well accepted prognostic factors for GIST, including tumor stage, tumor location, mitotic index (defined as mitoses per 5 mm^2^ on histopathologic analysis), and treatment status. Even when controlling for these variables, the *M2C molecular subtype* was a significant predictor of EFS (HR 26.34, 95% CI 2.6 – 268.4). (**Figure 5D**). Finally, we subset analyzed 45 *KIT* exon 11 mutant GIST lacking secondary *KIT* mutations **(Supplemental Figure 5C)**. Clinically, we would assume these tumors all to be molecularly similar independent of tumor location. While more than half (N=23) were IM2, the remaining 22 tumors included the other 5 molecular subtypes [range: 2-7 tumors (4%-16%)]. Overall, 16% (n=7) had events. When we analyzed the corresponding molecular subtypes, 43% of M2C (3/7), 33% of M1 (2/6), and 9% of IM2 (2/23) tumors were linked to events. Taken together, these findings suggest that transcriptional state heterogeneity can add to our understanding of homogenous genomic driver alterations to stratify patients.

We then externally validated our *molecular subtype* findings by analyzing 75 GIST samples collected by the Life Raft Group, an international GIST patient advocacy organization. These tumors underwent bulk RNA sequencing and were deconvolved into *cell state* abundances **(Supplemental Figure 6B)** with 44 (58.6%) samples closely matched to at least one of our *molecular subtype* profiles (*i.e.,* had an IC >0.50). An additional 9 (12%) samples were just below the matching threshold (IC 0.4 - 0.49) **(Supplemental Figure 6C)**. Like our test cohort, associations were noted between *molecular subtypes* and tumor locations, as well as mutational profiles. Again, *IM2* tumors were associated with *KIT* mutant gastric tumors (p < 0.05), while *M1* tumors were primarily small bowel GIST (p < 0.01).

Returning to our earlier insights into tumor cell evolution *in vitro* with treatment (**Fig. 4 & Supplemental Fig. 3**), we conducted a small pilot analysis of patients who underwent multiple rounds of sequencing during their care. Three of four patients received treatment throughout their course, though one patient was untreated during the entirety of sampling. Two patients had small bowel GIST (Patients 1 and 2), one had gastric GIST (Patient 3), and one patient was sampled after previously being treated for recurrent, metastatic GIST (Patient 4). We performed deconvolution as described above and matched the resultant cell state abundances with our molecular subtypes (**Supplemental Figure 7A-B**). The untreated patient (Patient 3) had minimal changes in cell state proportions. With treatment, Patient 1 changed from an *M1S* subtype to an *M1* subtype. While the molecular subtype did not change in two patients, we noted emergence of an M2 cell state in Patient 2 and a reduction in the Cycling cell state in Patient 4 **(Supplemental Figure 7C)**. In summary, our deconvolution approach can identify changes in cell state composition with treatment and tumor evolution.

Finally, utilizing single sample Gene Set Enrichment Analysis (ssGSEA) we sought to determine the enriched gene sets and pathways in these *molecular subtypes* (**Supplemental Figure 8, Supplemental Data)** {Barbie, 2009 #82}. The *IM2* subtype, which is associated with the ICC cell state, was associated with enrichment in pathways including ERBB3, PIK3CA/ATK/mTOR and Shh. Meanwhile the more aggressive *M2C* subtype was associated with pathways including BER, PolK, and E2F. The *M1S* subtype, associated with SDH-deficient tumors, was enriched in pathways involved with ER stress. This analysis represents putative molecular pathways that can be investigated for targeting these molecular subtypes in the future.

## DISCUSSION

In this study, we performed pooled analyses of single-cell RNA sequencing (scRNAseq) of human gut cells using publicly available datasets and created the largest reported reference map of the GI tract mesenchyme and enteric nervous system. Furthermore, to interrogate the tumor cell heterogeneity of human GIST, we performed single-nuclei RNA sequencing (snRNAseq) of 16 primary GIST, also representing the largest such study to date. We then performed reference mapping of GIST tumor cells to the gut reference map to identify distinct tumor *cell states*. No tumors were homogenous, and none were compromised of a single *cell state*. Thus, every tumor had variable intra-tumoral heterogeneity. Our findings also suggest that two distinct Mesodermal lineages may give rise to cells that can transform into GIST. Moreover, more than one-third of tumor cells mapped to fibroblastic (i.e., *PDGFRA* mutant) or smooth muscle cell (i.e., *SDH* mutant) expression profiles, while less than two-thirds of tumor cells mapped to ICCs (postulated precursor cells of GIST). We then utilized the presence of one dominant *cell state* or the combination of two dominant *cell states* to investigate the intra-tumoral and inter-tumoral heterogeneity of GIST by categorizing clinical samples into six different *molecular subtypes* that were associated with clinicopathologic features (e.g., tumor location and mutation status) and/or patient outcomes (i.e. event-free survival). Finally, we investigated how treatment is associated with transient changes in tumor cell state and composition. Thus, we present a novel approach for utilizing single cell transcriptomics to define common *cell states*, which we then utilize to define tumor cell heterogeneity of GIST through identifying 6 unique *molecular subtypes* of GIST. Taken together, we begin to unravel the complexity of each tumor by quantifying intra- and inter-tumoral heterogeneity that has biologic and prognostic significance.

This study highlights the fact that GISTs are without question heterogeneous beyond their respective mutation profiles. Previous studies have characterized other tumor *molecular subtypes*, including in glioblastoma^54^ and pancreatic cancer^55^. Thus, tumor heterogeneity is not unique to GIST. However, while bulk sequencing has previously identified the coarse-grained subtypes, single cell sequencing and deconvolution of bulk transcriptomes can be leveraged, as our study now suggests, to characterize GIST by functional *molecular subtypes*. By using *molecular subtypes* to further characterize tumors as a mixture of one or two more dominant *cell states*, we can both capture the heterogeneity of these tumors and highlight that tumors do not exist in single tumor *cell states*. Moreover, we provide an easy nomenclature that indicates tumor heterogeneity, as well as identifies which *cell states* may need to be targeted with therapies to eradicate both the majority and minority tumor cell types^56^.

While we have determined that 6 *cell states* exist, we also note that only 6 predominant *molecular subtypes* (i.e., combinations of these 6 cell states) tend to occur in our samples. This suggests a process of “functional convergence” exists by which GIST evolve to a limited set of viable configurations that manifest as our 6 *molecular subtypes*. Furthermore, certain *molecular subtypes* seem to have unique characteristics and tumor biology. For instance, we demonstrated that tumors within the distal stomach had higher percentages of non-ICC *cell states* (i.e., fibroblast-like and SMC-like). In turn, the *molecular subtypes* were also associated with particular locations, such as the IM2 subtype being mostly associated with the stomach and improved EFS, whereas the M1 subtype was associated with the small bowel. In regard to tumor biology, we noted that subtypes enriched for specific functional pathways, such as ERBB3, PIK3CA/ATK/mTOR or ER stress pathways. However, further studies need to be conducted to develop a more fundamental understanding of the biologic differences between these subtypes.

Despite advances in therapeutic options for GIST, many patients continue to succumb to this disease because they eventually develop TKI resistance. While resistance to imatinib and other TKIs have been identified through secondary mutations in *KIT*, for some patients’ resistance to therapy may develop without the presence of these mutations^57^. Another important clinical issue is that the natural course of the disease varies among patients. In some patients, GISTs are an indolent, slowly progressive disease, while in others, GISTs spread aggressively and are ultimately fatal despite sharing the same mutational profile. Studies have demonstrated that certain clinical and pathological attributes of GIST can be used to separate localized tumors into higher risk versus lower risk of recurrence following resection, including tumor size, mitotic index, and tumor location ^58–60^. These factors dictate the use of adjuvant therapy for patients that might be at high risk of recurrence. Interestingly, we found that tumor location and mitotic index were less significant predictors of EFS than the *molecular subtype* classification. The *molecular subtype* proved to be the best predictor of worsened outcomes, and likely reflects the fact that tumor location, mitotic index, and size of a tumor may all be indirect measures of a tumor’s *cell state* composition, and that a tumor’s *molecular subtype* captures all these clinical attributes.

Other studies have also attempted to use expression levels of specific genes for prognostication^42^. While our study confirms some of these previous findings, our study supports the fact that a single gene or protein (i.e., *BARX1* or *HAND1*) cannot be used alone for GIST prognostication. From our study, we note that 50% of the patients in our cohort had the *IM2 molecular subtype*, which would classically be responsive to imatinib. In addition, in our pilot study examining tumor compositions being treated over time, we noted that all four of these patients had *KIT* exon 11 mutations. Despite this, each patient had significantly different tumor compositions when interrogated by cell state and *molecular subtype*, with varying degrees of response. Thus, the molecular subtypes, which consider the absolute and relative expression of large gene sets, are required to fully capture the biology of GIST. They provide a novel and perhaps better predictor of outcomes worthy of further investigation in the future. Ultimately, by characterizing tumors by *molecular subtype*, our findings may allow for better prognostication, and more personalized treatment algorithms for GIST patients.

The current study has several limitations that should be considered when interpreting our findings. First, with GIST being a rare disease, the sample size of snRNAseq samples is small, with only 16 samples. Second, these samples are quite heterogenous in location, treatment status, and mutation status. We favored a sample group that was comprehensive of all possible *cell states* and subtypes, but further studies with larger sample sizes and more replicates of each specific mutational subtype are needed to corroborate our findings. Third, these samples only represent primary tumors and not metastases. Thus, there may have been a bias that more indolent disease is overrepresented in our dataset. Studies with snRNAseq on metastatic samples should be performed to examine if there are differences between primary and metastatic tumors in the same patient. Fourth, it is important to note that large GIST or pretreated GIST very frequently have areas of necrosis as well as different states of proliferation. Therefore, single samples may not capture all aspects of tumor heterogeneity and/or tumor biology. Fifth, our model was limited to one major institution and a small validation cohort of international patient tumors. Finally, only 58.6% of our validation cohort met the IC threshold for matching our established molecular subtypes, indicating that these subtypes must be tested and refined on a larger number of patient samples to be properly clinically validated.

In conclusion, our major findings and their implications for understanding GIST biology are: 1) that GIST are composed of non-ICC *cell states* therefore raising the possibility that non-ICC mesodermal cells may transform into GIST; 2) that combinations of 6 identified *cell states* define 6 specific *molecular subtypes* that are associated with particular driver mutations, tumor locations, and outcomes; and 3) these findings underscore the importance of utilizing transcriptomics to refine management strategies and improve patient prognostication.

## METHODS

### Tissue Isolation

Human GIST tissue was acquired with Institutional Review Board (IRB) approval from the University of California San Diego (IRB# 181755) using the UC San Diego Biorepository and Tissue Technology Shared Resources Center (La Jolla, CA). All patients provided written informed consent for the use of their deidentified biospecimens for research purposes. Resected human GIST were maintained on ice until arrival at the biobank. A pathologist or pathology physician assistant selected regions of tumor away from the margins that were deemed representative of the tumor specimen. These specimens were snap frozen at −80 °C until sequencing was performed.

### Single-nucleus RNA-seq data generation

Single nucleus RNA-seq was performed at the Center for Epigenomics at UC San Diego using the Droplet-based Chromium Single-Cell 3’ solution (10x Genomics). Briefly, nuclei were isolated from frozen tumor tissue using a gentleMACS dissociation protocol (Miltenyi). Approximately 20-50 mg of frozen tumor tissue was suspended in 2 ml of MACS dissociation buffer consisting of 5 mM CaCl2 (G-Biosciences, R040), 2 mM EDTA (Invitrogen, 15575-038), 1X protease inhibitor (Roche, 05-892-970-001), 3 mM MgAc (Grow Cells, MRGF-B40), 10 mM Tris-HCl pH 8 (Invitrogen, 15568-075), 0.6 mM DTT (Sigma-Aldrich, D9779), and 1 U/µl of RNase inhibitor (Promega, N2111B) in water (Corning, 46-000-CV) and placed on wet ice. Next, samples were homogenized using gentleMACS dissociator (Miltenyi) with gentleMACS M tubes (Miltenyi, 130-096-335) and the “protein_01” protocol. The suspension was filtered through a 30 µM CellTrics filter (Sysmex, 04-0042-2316). The M tube and filter were washed with 1 ml of MACS dissociation buffer and combined with the suspension, centrifuged in a swinging bucket centrifuge (Eppendorf, 5920R) at 500 rcf for 5 minutes (4°C, ramp speed 3/3). Supernatant was removed and pellet was resuspended in 500 µL of lysis buffer consisting of 0.1% Triton X-100 (Sigma, T8787-100ML), 1X protease inhibitor (Sigma, NC0969110), 1 mM DTT (Sigma, D9779), and 1 U/µL RNase inhibitor (Promega, N2111B) in 2% BSA (Sigma, SRE0036) in PBS (Corning, 21-040-CV) and incubated on a rotator for 5 minutes at 4°C. Lysate was then centrifuged at 500 rcf for 5 minutes (4°C, ramp speed 3/3) and the supernatant was removed. Pellet was resuspended in 400 µL of sort buffer consisting of 1 mM EDTA (Invitrogen, 15575020) and 1 U/µL RNase inhibitor (Promega, N2111B) in 2% BSA (Sigma-Aldrich, SRE0036) in PBS (Corning, 21-040-CV) and stained with DRAQ7 at a final concentration of 3 µM (Cell Signaling, 7406) on ice for 10 minutes.

Nuclei were sorted using a SH800 sorter (Sony) into 50 µL of collection buffer consisting of 1 U/µL RNase inhibitor (Promega N211B) in 5% BSA (Invitrogen, AM2616/AM2618); the FACS gating strategy isolated and sorted nuclei based on particle size and DRAQ7 fluorescence. Sorted nuclei were then centrifuged at 1000 rcf for 15 min (4°C, run speed 3/3) and the supernatant was removed. Nuclei were resuspended in 35 µL of reaction buffer consisting of 1 U/µL RNase inhibitor (Promega, N2111B), and 1% BSA (Invitrogen, AM2616/AM2618) in PBS (Corning, 21-040-CV) and an aliquot was used for counting on a hemocytometer by staining with Trypan Blue (Life Technologies, 15250-061).

Nuclei were then loaded onto a Chromium Controller (10x Genomics) to perform single-nucleus profiling of gene expression using the Chromium Next GEM Single Cell 3’ GEM Kit v3.1 (10x Genomics, 1000123) and the Chromium Next GEM Chip G Single Cell Kit (10x Genomics, 1000120). Reverse transcription on barcoded transcripts and CDNA amplification for 12 cycles were performed using a T100 Thermocycler (BioRad). CDNA cleanup was performed using SPRISelect reagent (Beckman Coulter, B23319) and library concentration was assessed using Qubit dsDNA HS Assay Kit (Thermo-Fischer Scientific).

Single-nucleus RNA-seq libraries were then generated using the Chromium Next GEM Single Cell 3′ Library Kit v3.1 (10x Genomics, 1000157) and a T100 Thermocycler (BioRad) to fragment CDNA and anneal Illumina sequencing adapters. Individual libraries were PCR amplified using index primers from the Dual Index TT Set A (10x Genomics, 1000215) according to manufacturer specifications. SPRISelect reagent (Beckman Coulter, B23319) was used for size selection and clean-up steps. Final library concentration was assessed by Qubit dsDNA HS Assay Kit (Thermo-Fischer Scientific, Q32854) and fragment size was checked using Tapestation High Sensitivity D1000 (Agilent) to ensure that fragment sizes were distributed normally about 500 bp. Libraries were sequenced using the NextSeq500 and a NovaSeq6000 (Illumina) with these read lengths: 28 + 10 + 10 + 91 (Read1 + Index1 + Index2 + Read2).

### Analysis of human single-nucleus RNA sequencing data

#### Preprocessing and Preliminary Quality Control

Cell Ranger^61^ was used to process the raw FASTQ data and generate cell by gene read count matrices. In the Cell Ranger pipeline, *‘cellranger count’* was used with all default parameter settings. Data analysis was primarily performed using Seurat^62–64^.Cells were removed that contained mitochondrial gene expression percentages more than 0.2%, as well as those cells containing less than 200 or greater than 3000 genes per cell. Additional quality control measures were performed using DoubletFinder^65^ and SoupX^66^ to remove additional heterotypic doublets and ambient RNA expression, respectively.

#### Normalization and Clustering

Mitochondrial and ribosomal genes were removed prior to normalization and integration. The filtered cell expression matrix was then normalized using the *LogNormalize* method in Seurat^62–64^. Variable genes were selected based on a residual variance threshold of 1.3. Principal Component Analysis (PCA) was then performed on the resulting scaled expression values. A shared-nearest-neighbor (SNN) graph was constructed using the Seurat FindNeighbors function with the “Annoy” method, using the first 20 principal components (PCs), a cosine distance metric, and number of nearest neighbors count set to 20. Clusters were identified from the SNN graph using the Seurat FindClusters function with the Louvain algorithm and resolution parameter set to 0.8. The UMAP values were calculated from the top 20 PCs using the uwot R package (version 0.1.8) with cosine distance metric and n.neighbors set to 20. For the cell-type-specific UMAP values, PCA was repeated within each subset and the top 30 PCs were used instead with the same parameters otherwise. The number of PCs were selected based on heuristic methods including generating an ElbowPlot, a ranking of principle components based on the percentage of variance explained by each one in order to inform how many PCs are required to capture the true signal of the data ^62–64^.

#### Harmony Batch Correction

Raw read counts were normalized with the Seurat R package using the log-normalization method with default parameters. The top 3000 most variable features were then selected using the variance stabilizing transformation method. The resulting subset was scaled and centered, and principal component analysis (PCA) was performed using the default parameters. To ameliorate sample or patient-specific batch effects, the Harmony R package (v0.1.0) was applied to the Seurat object using the grouping variables patient ID and batch^67^. The resulting Harmony embedding was used to perform UMAP dimensional reduction, neighbor finding, and cluster finding with the first 30 dimensions and resolution of 0.5.

#### Annotation of Cell Clusters and Identification of Tumor Cells

Marker genes for all cell clusters were computed using the FindAllMarkers function from Seurat for each cluster, with a log-fold change threshold of 0.3. Only genes expressed in a minimum of 50% of all cells in a cluster were used. Cell clusters were identified as tumor cells based on expression of *KIT*, *ANO1*, and *ETV1*. Clusters expressing all three of these markers were labeled as *KIT*^+^ tumor cells. However, given the small prevalence of *KIT*^-^ GIST, cells with *ANO1* expression were tentatively labeled as *KIT^-^* GIST. To ensure that these cells were not pericytes, which are also known to express *ANO1*, we also checked that pericyte markers *RGS5*, *ANGPT1*, and *NOTCH3* were not concurrently expressed. Finally, we created a GIST composite score consisting of multiple GIST markers (*KIT, ANO1, ETV1, PRKCQ*, *PDE1A*, *PROM1* (CD133), *FOXF1*) using the AddModuleScore function in Seurat ^68^.Populations with high GIST composite scores were considered tumor cells. This function calculates the average expression levels of each set of genes on a single cell level, subtracted by the aggregated expression of control feature sets. All analyzed features are binned based on averaged expression, and the control features are randomly selected from each bin.

For other non-tumor cells, using the markers obtained earlier in the FindAllMarkers function, we performed annotations using common marker databases including CellMarker^25^ and PanglaoDB^69^ to identify possible cell types. We confirmed these annotations by performing comprehensive literature review.

#### Reference Map Creation

In order to create the reference map, single cell and single nuclei data was collected from 19 studies encompassing more than 700 samples and over 1,000,000 cells^20–39^. CellxGene was primarily used to download the data^70^. Reference mapping was performed using the Symphony package. Mitochondrial and ribosomal genes were removed from all datasets prior to normalization and integration. In addition, all datasets were annotated to include technical aspects of each (i.e., single cell vs single nuclei), patient ID, and batch. The datasets were then merged as follows. Scaling, PCA, and normalization were performed as before. Harmony batch correction was performed on the four datasets, this time accounting for batch, patient ID, and sequencing technology for integration. Following integration, clustering and projection using UMAP were performed as described above. The Symphony package was then used to create a low dimensional embedding of the reference map.

#### Projection onto the Reference Map

Tumor cells were isolated as described above. Next, projection of the tumor cells was performed using the Symphony package. Using the MapQuery function, tumor cells were projected onto the reference map based on low dimensional embeddings. The MapQuery function subsets the query expression data by the same variable genes used in reference building and scales the normalized expression of the genes by the same means, Next, the Symphony k-nearest neighbors (KNN) algorithm was then used to perform label transfer from the reference dataset to the GIST query dataset. These annotations were then saved.

#### Tumor Subtype Characterization using ssGSEA

We identified enriched pathways in each tumor subtype by performing single sample Gene Set Enrichment Analysis (ssGSEA). Pathway databases were obtained from MSigDB and included Hallmark pathways (H), canonical pathways (C2), and gene ontology pathways (C5). This ssGSEA analysis was run with default parameters, yielding a set of enrichment scores for each sample. A two-sample t-test was performed between the samples associated with each tumor subtype and all other samples to identify the relative enrichment of pathways in each tumor subtype. The top 200 pathways ranked by t-statistic were chosen for each subtype and a heatmap of the scaled enrichment scores for the selected pathways was generated.

#### Deconvolution and Validation Cohorts

Bulk RNA sequencing was obtained via Tempus as FASTQ files. Transcript abundances were obtained using kallisto^71^, using paired mode under default settings, using the latest index files based on Ensembl v96 transcriptomes (https://bit.ly/3ytr4OL). Data from Life Raft Group included RNA sequencing data provided courtesy of Columbia University and were single read FASTQ files.d therefore abundances were obtained using kallisto using the options ‘--single’, ‘-l 100’, ‘-s 20’, ‘-b 2, and’ ‘-t 2’. Counts and normalized TPM counts were obtained using the *tximport* package^72^ using the function ‘tximport’, with the settings type = “kallisto” and txOut = TRUE.

Deconvolution was performed using the BayesPrism software package^50^. The software uses an Expectation-Maximization (EM) approach to estimate the distribution of cell-type composition and cell type-specific gene expression from bulk RNA-seq expression of tumor samples^73^. Using single nuclei data, cells were assigned to 6 different states based on reference mapping described above. The parameters cell.type.labels and cell.type.states were treated as equivalent. Gene filtering of the single cell reference was performed under recommended settings, removing non-protein coding genes only, and removing mitochondrial, ribosomal, ChrX and ChrY genes. Genes with zero or low expression (genes with less than 3 counts among all cells) were also excluded. Finally, to improve the accuracy of the deconvolution, and given that transcriptional profiles are similar between *molecular subtypes*, we subsetted on signature genes only using the get.exp.stat function under 10x recommended settings (pseudocount = 0.1). A Prism object, with contains all data required for running BayesPrism, namely, a scRNA-seq reference matrix, the cell type and *cell state* labels of each row of reference, and the mixture matrix for bulk RNA-seq, was created using input.type = ‘count_matrix’ and key = NULL under recommended settings. Deconvolution was then performed on the Prism object using ‘run.prism’ to obtain final tumor composition.

#### Defining Molecular Subtypes and Matching Datasets

In order to define *molecular subtypes*, we clustered 75 of the deconvolved samples using a Self-Organizing Map (SOM) and used *6 subtypes* as these captured most of the variation seen in those samples^74^. For each subtype, we averaged the *transcriptional profile signature* of the corresponding member samples to define characteristic *molecular subtype profiles*. In this way, these *molecular subtypes* help define groups of samples with similar *transcriptional profile abundances* and representation of *cell states*. In order to map an independent dataset onto the *molecular subtypes*, e.g., the Columbia samples, we match each independent dataset’s *transcriptional profile abundances* against the 6 *molecular subtype profiles* and determine the best match according to the Information Coefficient (IC) ^51–53^. The IC is a measure of correlation similar to the Pearson correlation coefficient but better at detecting non-linear association. The IC values are in the range [-1, 1] with 1 being perfect correlation and −1 perfect anti-correlation, and zero for no correlation. For matching independent samples, we set up an IC threshold of 0.6 so the samples have to match with an IC above that threshold in order to be considered as “matching” a *molecular subtype.* For the second cohort 43 out of 75 samples (57.3%) match with an IC higher than 0.6.

#### Statistical Analysis

Where appropriate, data are expressed as mean values plus or minus one standard deviation, and were compared when normality was approximated. Statistical analyses were performed using two-sided Wilcoxon Rank Sum tests for continuous data and Chi squared tests for categorical data, unless otherwise stated. Univariate Kaplan Meier tests were performed using a log-rank test to test for significance. Multivariate survival analysis was performed using Cox proportional hazards models. In cases where multiple comparisons were made, *P* values were adjusted using a Bonferroni correction.

## Acknowledgements

We appreciate funding support from the NIH T32 CA121938 Cancer Therapeutics (CT2) Training Fellowship (A.K. Sharma, M. Antkowiak); NCI Award F31CA257344 (A.T. Wenzel); NIH U01 CA27847 (J.K. Sicklick); NIH U01 CA184898 (J.P. Mesirov), NIH U24 CA220341 (J.P. Mesirov and P. Tamayo), NIH U24 CA248457 (J.P. Mesirov and P. Tamayo); NIH R01 CA226803 (J.K. Sicklick, P. Tamayo), FDA R01 FD006334 (J.K. Sicklick), NIH U01 CA278470 (J.K. Sicklick), Lighting the Path Forward for GIST Cancer Research (J.K. Sicklick), and GIST Support International (J.K. Sicklick). We also appreciate support from shared resources (GCBSR) at the UCSD Moores Cancer Center NIH P30 CA023100. This work was supported by the NCI Office of Cancer Target Discovery and Development (CTD^2^) award U01 CA 272610 (A. Califano), the NCI Outstanding Investigator award R35 CA 197745 (A. Califano), and the NIH Shared Instrumentation Grants S10 OD012351, S10 OD021764 and S10OD032433 (A. Califano).

## Declaration of Interests

J. Sicklick receives consultant fees from Deciphera, Aadi and Grand Rounds; serves as a consultant for CureMatch, received speakers’ fees from Deciphera, La-Hoffman Roche, Foundation Medicine, Kura, Merck, QED, SpringWorks and Daiichi Sankyo; and owns stock in Personalis and CureMatch.

Dr. A. Califano is a founder, equity holder, and consultant of DarwinHealth Inc., a company that has licensed propriety systems biology algorithms. Columbia University is also an equity holder in DarwinHealth Inc.

The Life Raft Group (S. Rothschild, D. Evans) has received program-related grants from Blueprint, Cogent Biosciences, Daichii Sankyo, Deciphera, Genentech, IDRx, Novartis, Pfizer, and Theseus.

## Data Availability Statement

Raw data for this study were generated at Center for Epigenomics at UC San Diego. Derived data supporting the findings of this study are available from the corresponding author upon request.

## SUPPLEMENTAL FIGURE LEGENDS

**SUPPLEMENTAL FIGURE 1. In-depth clustering analysis of the Human Gut Reference Map and snRNAseq Human GIST Atlas. A.** Markers calculated by differential expression between clusters of the Human Gut Reference Map. Dot size indicated percentage of cells within a given cluster expressing the gene of interest, and color indicates average expression. Clusters were previously defined in prior studies or datasets. Only 3 genes per cluster are shown for brevity. **B.** Left: Top markers based on differential expression for Seurat-defined tumor clusters in all *KIT*^+^ and *KIT^-^* tumor cells. Dot size indicated percentage of cells within a given cluster expressing the gene of interest, and color indicates average expression. Only 5 genes per cluster are shown for brevity. Right: Distribution of *KIT^+^* and *KIT^-^* tumor cells by cluster. **C.** Differentially expressed genes in treated versus untreated *KIT*^+^ and *KIT^-^* tumor cells. Dot size indicated percentage of cells within a given cluster expressing the gene of interest, and color indicates average expression. Only 5 genes per cluster are shown for brevity. **D.** Expression of common GIST markers in untreated (colored blue) and treated (red) samples.

**SUPPLEMENTAL FIGURE 2. Reference Mapping of Tumor Cells onto the Human Gut Reference Map. A.** Comparison of single cell and single nuclei RNA sequencing. Single cell RNA sequencing of two human GIST adapted from Mao *et al.* Cancer Sci. 2021, with annotated cell clusters ^45^. **B.** Single nuclei data from 16 human GIST were projected onto A, with good concordance. **C.** Heatmap indicating proportion of cells assigned to each *cell state* based on sample. Red indicates higher percentage of cells within that particular cluster. Sample mutation status and location data are displayed on the right. **D.** Tumor cells are projected onto the Gut Reference Map (grey scale). Tumor cells are colored by treatment status. Blue = Untreated, Red = Treated.

**SUPPLEMENTAL FIGURE 3. Pseudotime Analysis of GIST cell lines using Monocle. A.** Histograms demonstrating three GIST cell lines, T1 untreated cell lines (red), T1 cells treated with imatinib (blue, T1-IM), and T1 cells resistant to imatinib (orange, T1-IM^R^) over assigned pseudotimes from Monocle. **B.** Assigned cell states for T1 untreated cell lines over pseudotime. The three most common cell states (ICC, Mesoderm_1, and SMC) are displayed, other cell states were omitted due to low number of cells assigned to these states. **C.** Histogram of assigned cell states for T1 cells treated with imatinib (T1-IM) over pseudotime. **D.** Histogram of assigned cell states for T1 cells resistant to imatinib (T1-IM^R^) over pseudotime.

**SUPPLEMENTAL FIGURE 4. Pseudomixture Generation and Validation of Deconvolution. A.** Two hundred pseudomixtures from snRNAseq data were generated randomly and ensured to include all 6 *cell states*. Right = Actual fractions calculated at the time of pseudomixture generation. Left = Predicted fractions based on BayesPrism decovolution method. Middle = Correlation between Predicted and Actual fractions. **B.** Overall distribution of correlations between actual and predicted fractions.

**SUPPLEMENTAL FIGURE 5. Molecular Subtype Analyses. A.** Molecular subtype labels of individual samples by general location (stomach, small bowel). **B.** Molecular subtype labels of individual samples by driver mutation. Each bar graph represents what proportion of samples in each location or mutation sub-group were assigned a molecular subtype. Only locations/driver mutations with n ≥ 4 were included. **C.** Molecular subtype distribution of N=45 patients with *KIT* exon 11 mutations without any secondary mutations.

**SUPPLEMENTAL FIGURE 6. Molecular Subtype Projections Reveal Hetereogeneity of GIST. A.** *Molecular subtypes* of n=71 deconvolved patients (UC San Diego dataset) based on exon status and treatment status (p < 0.05, blue box, p<0.01, gold box). **B.** Deconvolution data of n=75 patients from Columbia University / Life Raft Group. Clinical metadata is shown below predicted abundances, including tumor mutation status, location, recurrence, and resistance. **C.** *Molecular subtype* projection of 75 patients. Forty-four patients (58.6%) met the IC threshold for matching. Associated metadata is included (p < 0.05, blue box, p<0.01, gold box).

**SUPPLEMENTAL FIGURE 7. Pilot Analysis of 4 Patients with GIST over time. A.** Four patients underwent various therapies over time and underwent multiple rounds of sequencing. Deconvolution of each sample demonstrating differences in tumor composition over time. Nomenclature: Patient *x.y* indicates Patient #*x* at relative timepoint *y*, with *y=*1 indicating the first time measurement, *y=*2 indicating the second time measurement, etc. **B.** Assignment of molecular subtypes over time for each patient **C.** Bar plot demonstrating change of composition over time for each patient. Treatment data and mutation status are noted for each patient.

**SUPPLEMENTAL FIGURE 8. Functional Enrichment Analysis of each Molecular Subtype.** Heatmap demonstrating top 200 enriched pathways by ssGSEA for each molecular subtype compared to all other molecular subtypes. Enrichment scores are normalized from scores 0 to 1.

**SUPPLEMENTAL DATA.** List of Top 200 enriched pathways by Molecular Subtype.

