## Supplemental Figures for "Single Nuclei-Derived Molecular Subtypes of Gastrointestinal Stromal Tumors Correlate with Clinicopathologic Features and Predict Clinical Outcomes"

#### Supplemental Figure 1

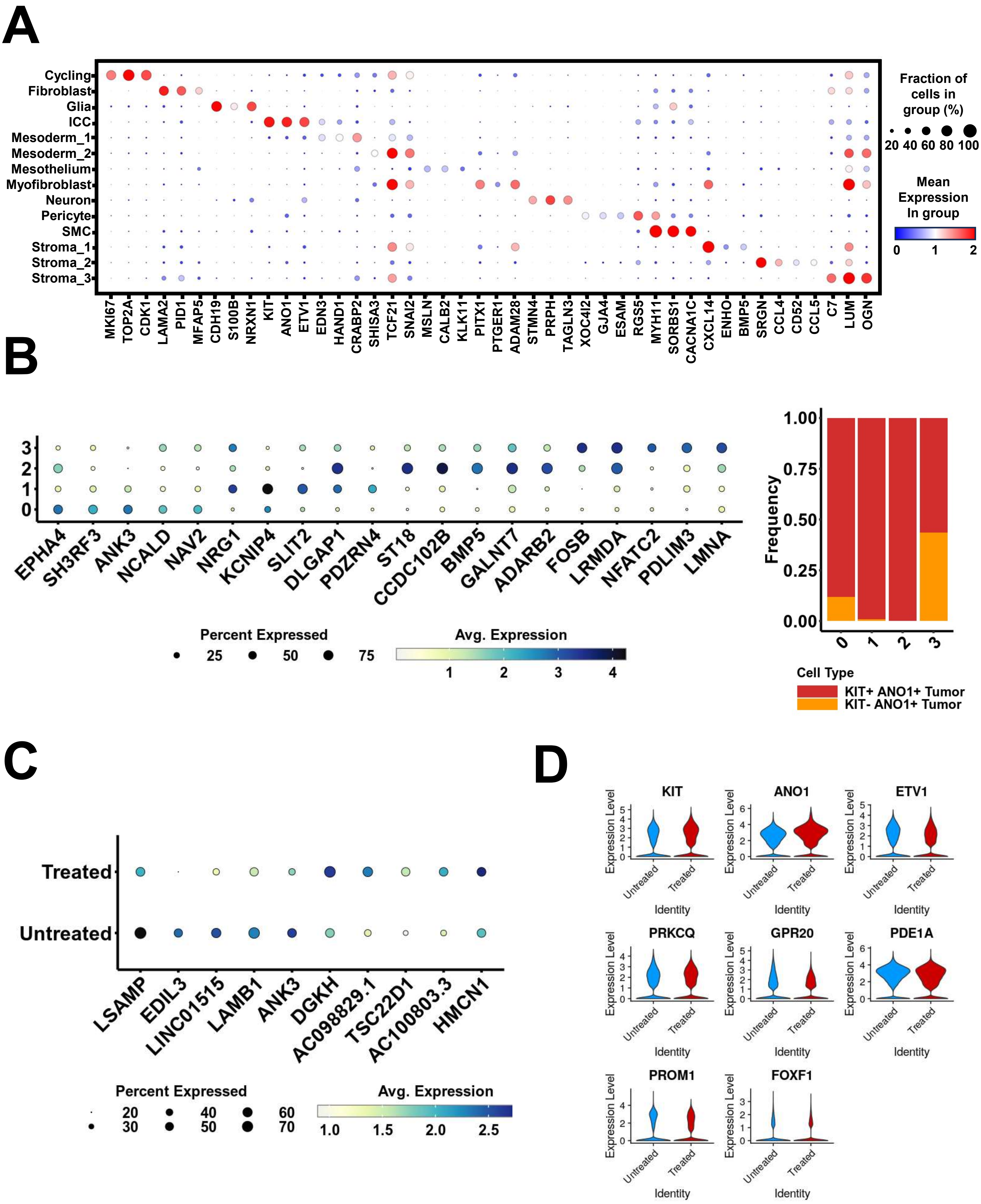

**Supplemental Figure 2**

**A**

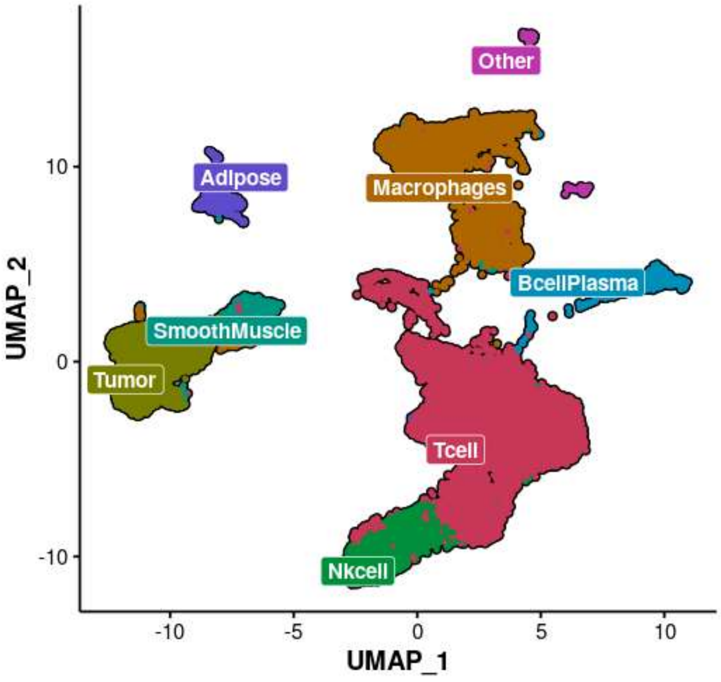


**Single Cell Reference of GIST (adapted from *Mao et al. Cancer Sci. 2021*)**

**B**

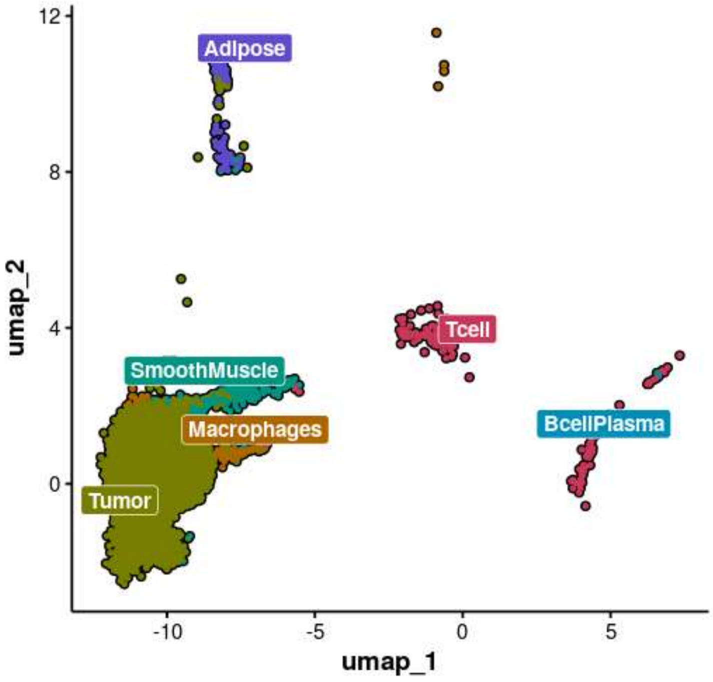


**Single Nuclei RNAseq data of tumor cell labeled by projection onto Panel A.**

**C**

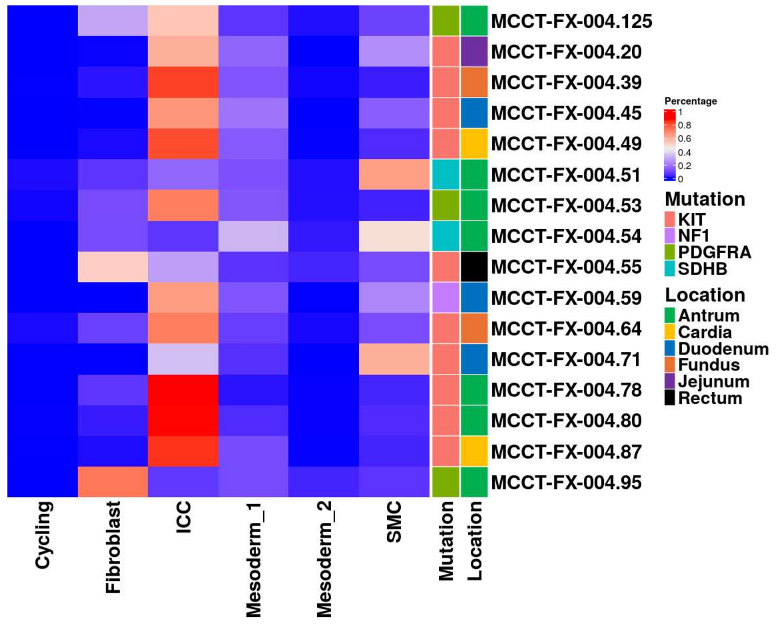


**D**

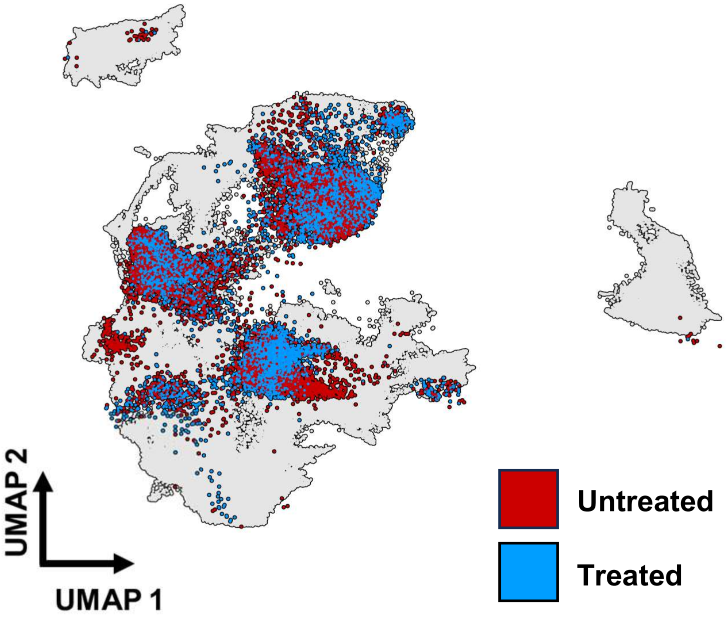

Supplemental Figure 3

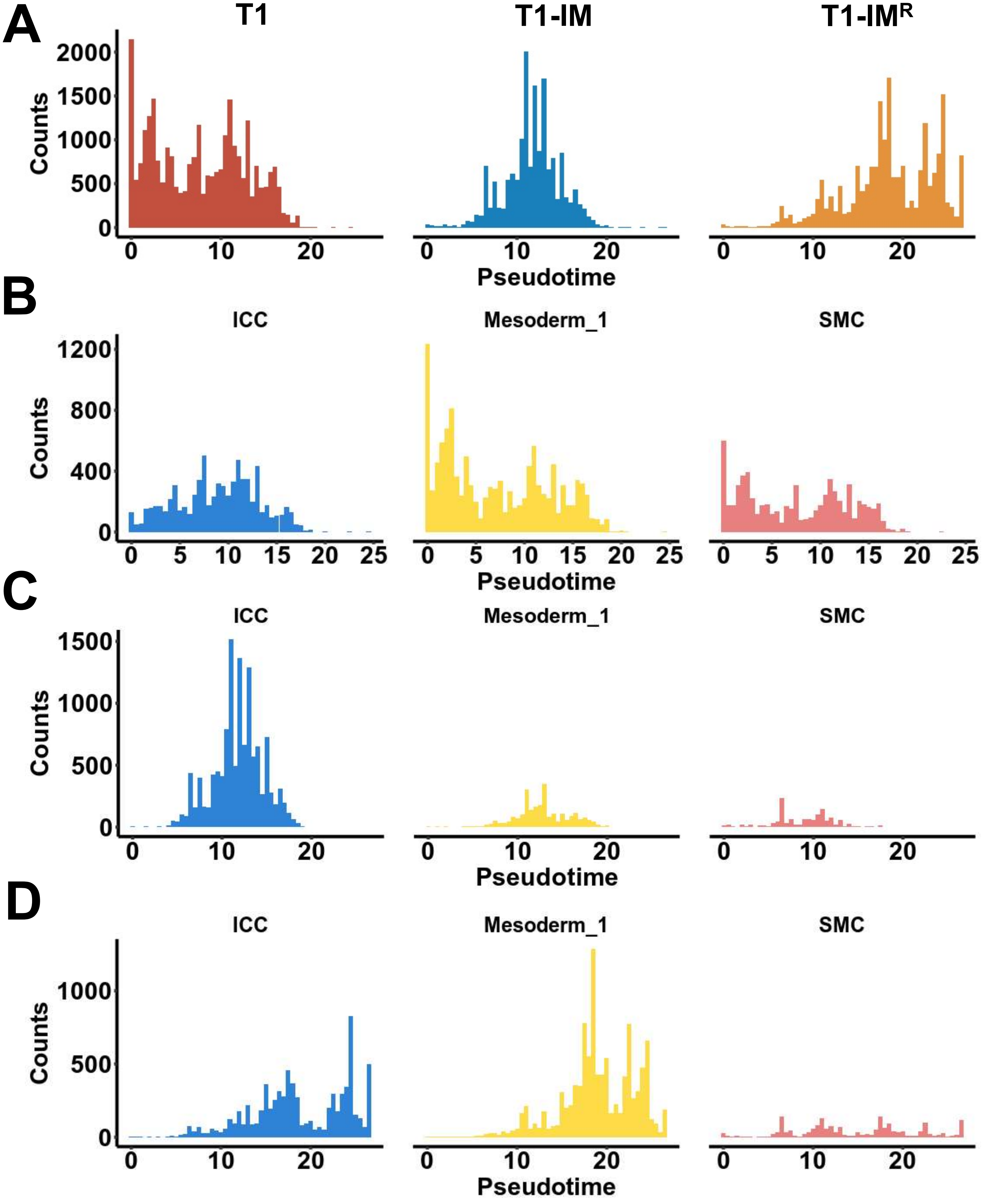

Supplemental Figure 4

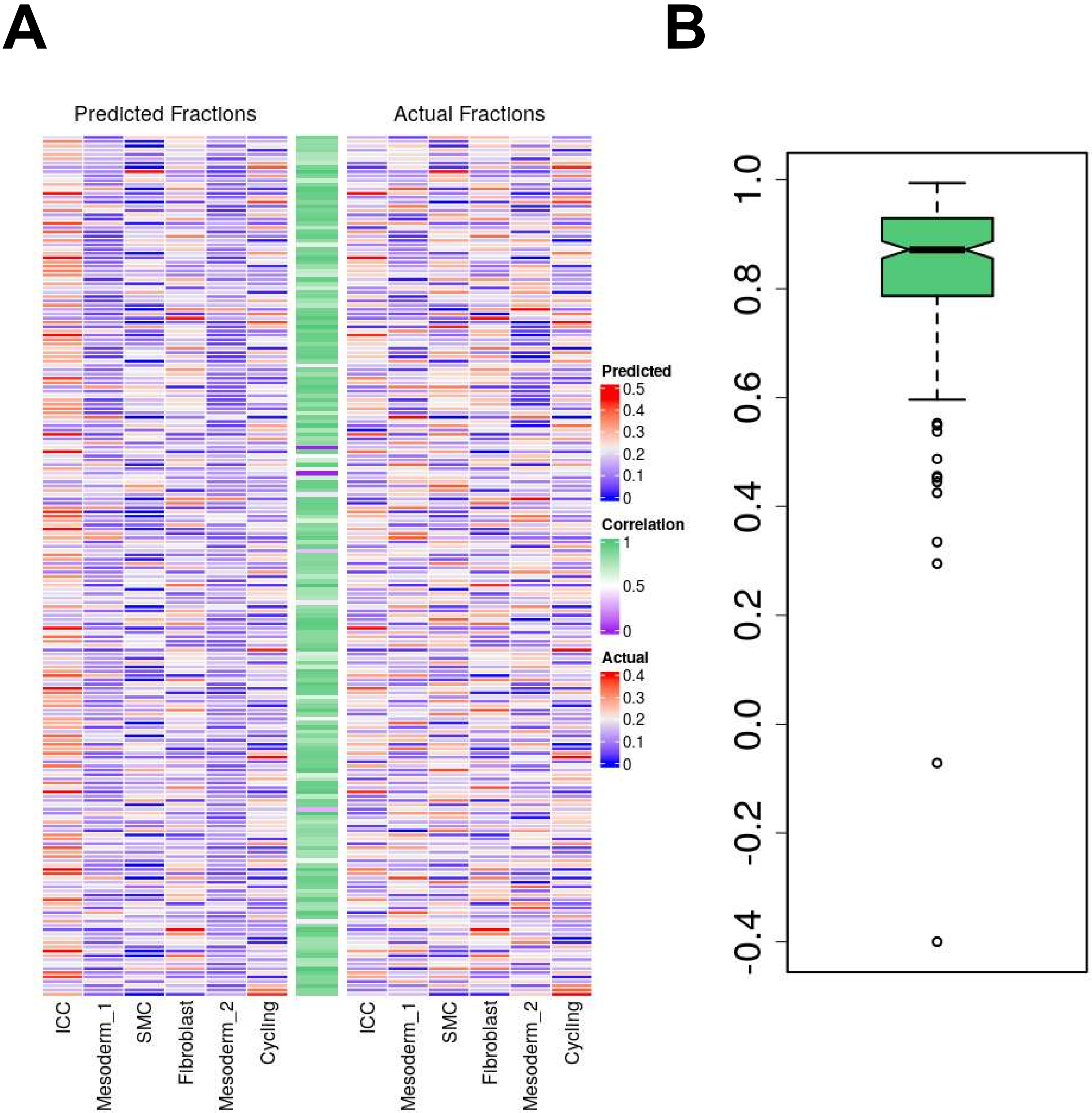

Supplemental Figure 5

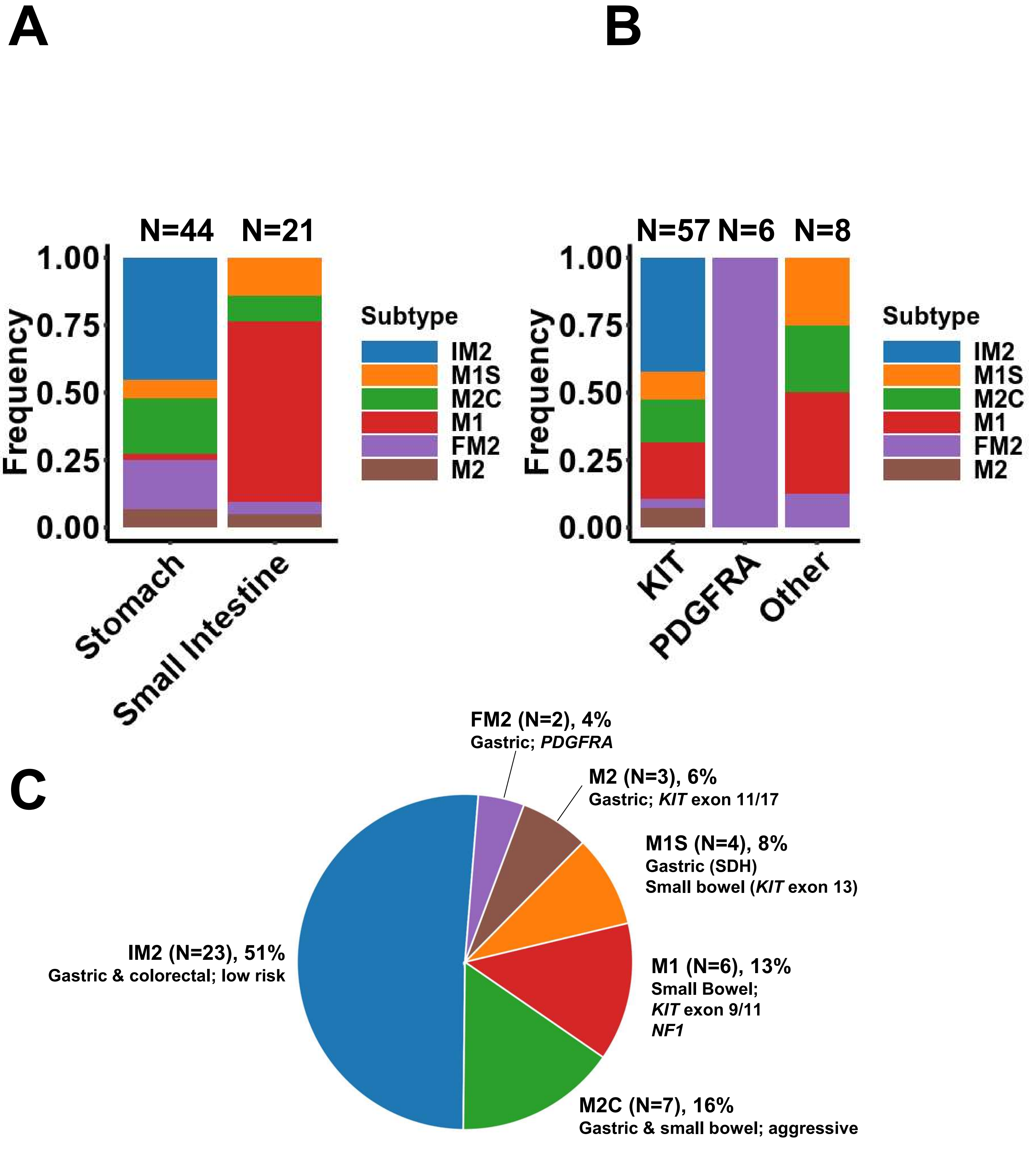

### Supplemental Figure 6

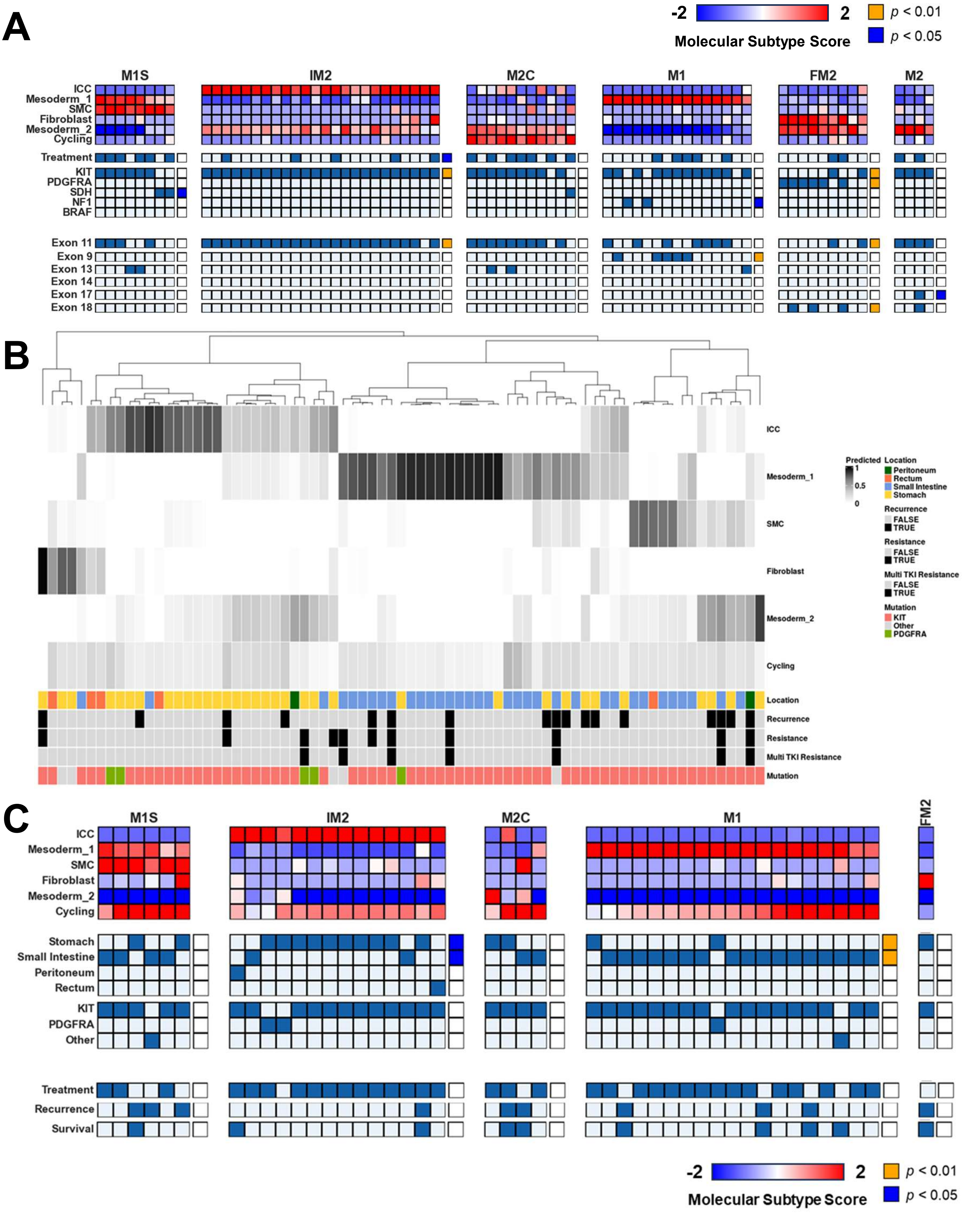

### Supplemental Figure 7

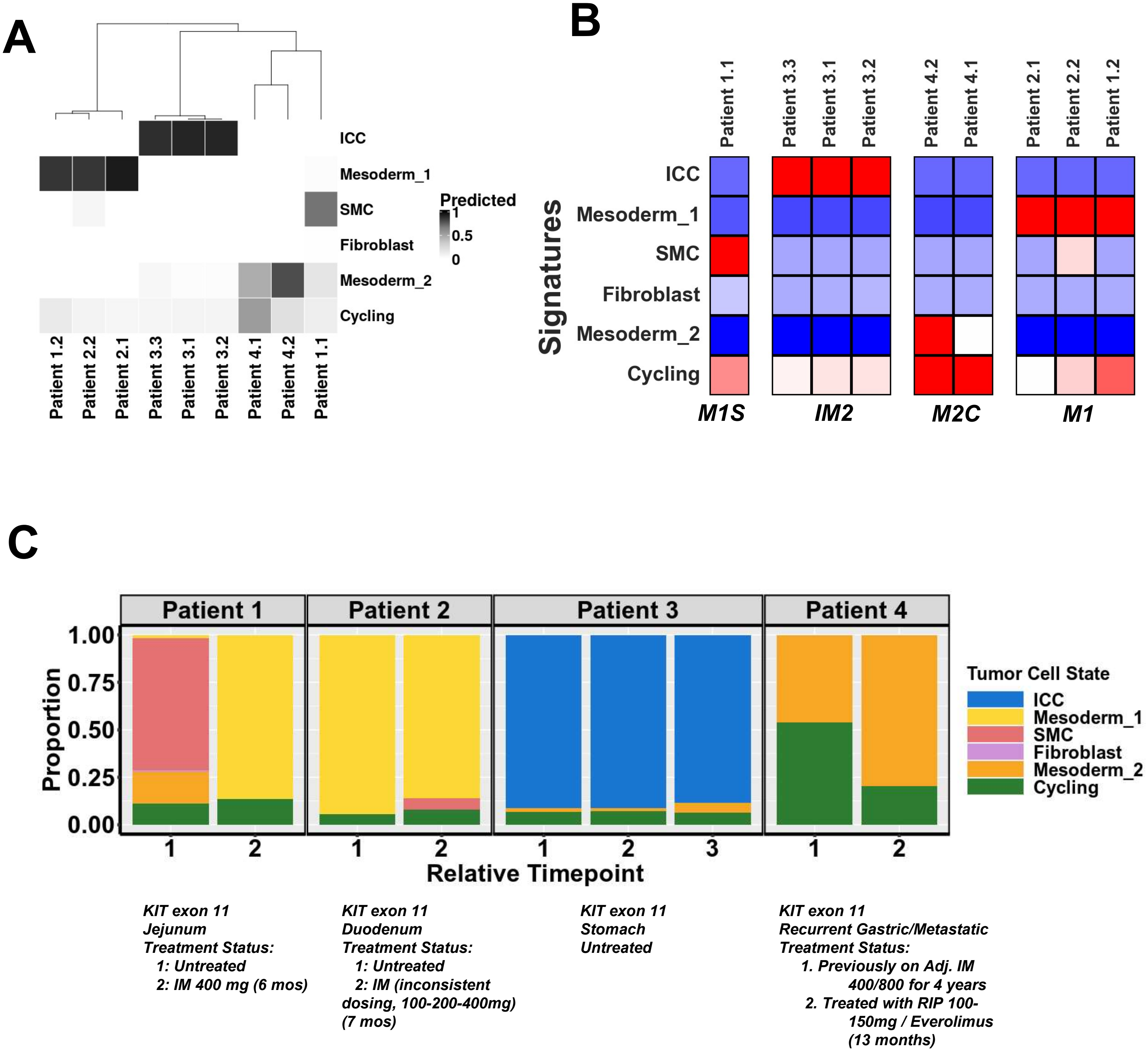

**Supplemental Figure 8**

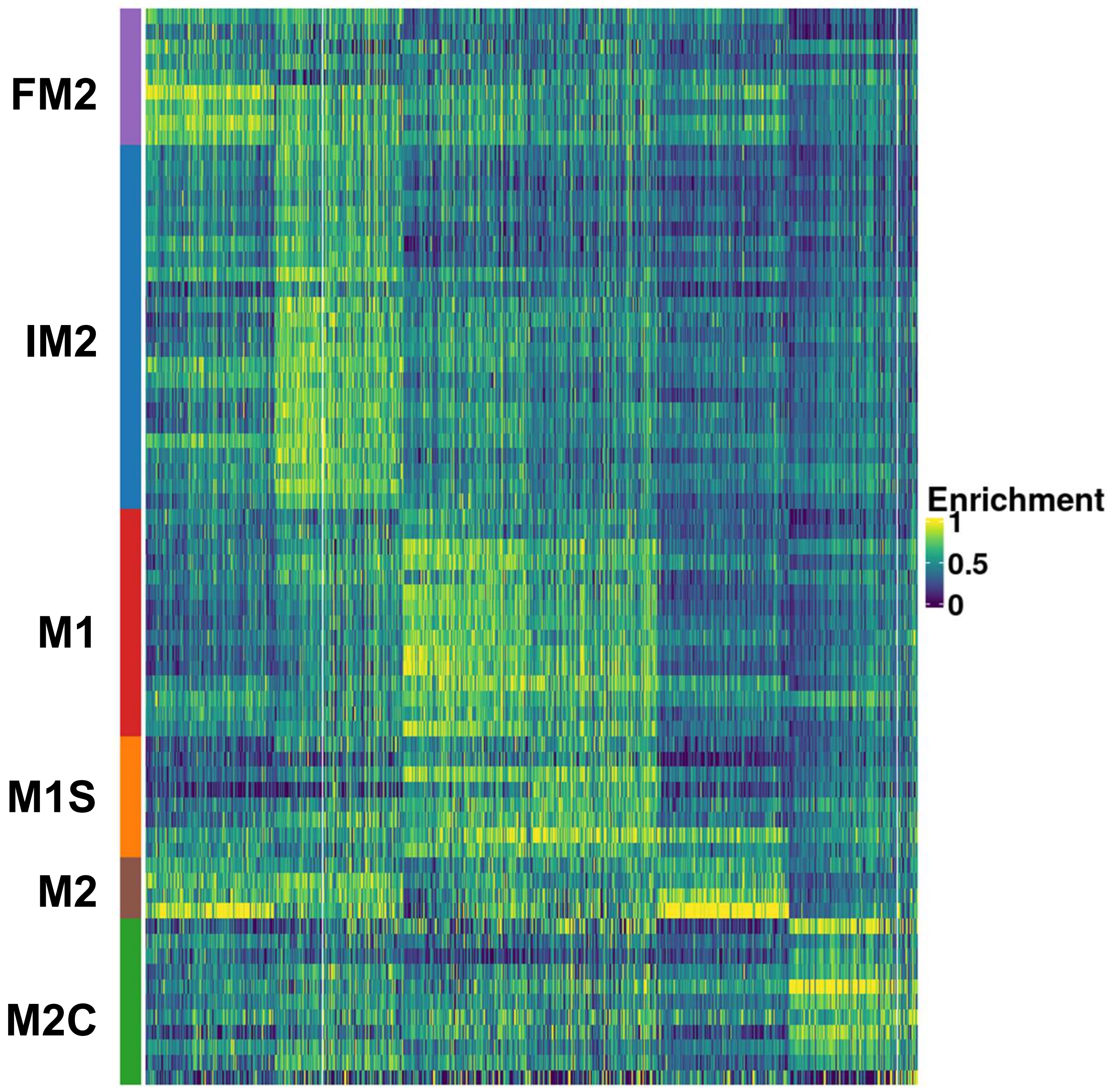
